# How Robust are Multispecies Coalescent Species Delimitations in Taxonomically Complex Systems? A Genomic Assessment Using Mediterranean *Tethya* Sponges

**DOI:** 10.64898/2026.07.04.735074

**Authors:** Joëlle van der Sprong, Frine Cardone, Sebastian Höhna, Simone Schätzle, Fabian Deister, Dirk Erpenbeck, Gert Wörheide, Sergio Vargas

## Abstract

Reliable species delimitation underpins biodiversity assessment but remains difficult for organisms with plastic morphology and few diagnostic characters. Multispecies coalescent (MSC) methods can delimit species from genomic data, yet they are rarely tested in taxonomically complex, marine invertebrate groups where they are arguably most needed. We used the three Mediterranean species of the genus *Tethya*, a rare, well-characterised system within the otherwise taxonomically difficult phylum Porifera—distinguished by multiple independent morphological and ecological characters—to evaluate how robust MSC-based delimitation is in such groups. Analysing 64 single-copy nuclear loci in BEAST2 and BPP, we compared constrained, hypothesis-testing approaches (BFD*, BFdriver, A10) with freer, heuristic ones (SPEEDEMON, A11), and examined their sensitivity to data type, clock model, priors, and the species-collapse threshold. All methods recovered the three recognised Mediterranean species, but the resolution of within-lineage structure was method-dependent. The hypothesis-testing approaches consistently supported six lineages, robustly across data types and model assumptions, whereas the heuristic approaches proved less stable. Configurations without *a priori* species hypotheses often failed to converge or were computationally intractable, a problem compounded by the relaxed clock. In SPEEDEMON the outcome changed with the collapse threshold. Because our system lacks an independent reference point to calibrate this threshold, any delimitation based on it is poorly constrained. We conclude that constrained, hypothesis-testing delimitation is the most robust and reproducible MSC approach, yielding a quantitative, model-based hypothesis that can be weighed against other lines of evidence to inform taxonomic decisions. By clarifying how these methods behave and how their outcomes should be interpreted, our study offers a practical guide for researchers working on comparably complex systems.

---

Species discovery and delimitation are fundamental to understanding both the patterns of biodiversity and the processes that drive it (Sites and Marshall 2003; Wiens 2007), which is critical in the context of ongoing global biodiversity loss (Cowie et al. 2022). Traditional approaches (i.e., alpha taxonomy) rely on descriptive morphology, with new species recognised when warranted by taxonomic authority. Although this works well for many organisms, it reaches its limits in groups that lack obvious diagnostic characters or in which morphology is plastic and difficult to interpret, complicating unambiguous species assignments (Adams et al. 2014; Bridge et al. 2024). DNA barcoding has since transformed species identification by using one or a few loci, often from the maternally inherited mitochondrial DNA, to delimit species from the ratio of intraspecific to interspecific variation: the barcoding gap (Hebert et al. 2003; Meyer and Paulay 2005). In practice, however, the extent of intraspecific variation is often poorly characterised, limiting the reliability of delimitation on this principle (Collins and Cruickshank 2013). Moreover, while single-marker methods are practical, they fail to capture the complexity of biological processes involved in ongoing speciation, such as ancestral polymorphisms, incomplete lineage sorting (ILS), or ongoing gene flow (Fujita et al. 2012; Xi et al. 2016; Adams et al. 2018). They often ignore phylogenetic uncertainty and treat divergence in a single gene as a direct proxy for species boundaries, which can lead to inaccurate or oversimplified species hypotheses (Degnan and Rosenberg 2009).

The advent of the multispecies coalescent (MSC) model has greatly improved species delimitation and discovery. By providing a probabilistic framework for modelling population histories and inferring species trees, the MSC accounts for gene tree-species tree discordance, which can arise from stochastic processes such as ILS (Rannala and Yang 2003; Fujita et al. 2012). It estimates population parameters and divergence times from multiple loci without assuming reciprocal monophyly, enabling quantitative inference of biologically complex scenarios. Nevertheless, the model has several limitations. First, the standard MSC assumes instantaneous speciation and no post-divergence gene flow, oversimplifying intraspecific dynamics and failing to capture hybridisation and introgression (Mallet 2005). Second, limited taxon sampling and geographic clines can confound population structure with speciation signal, assigning questionable species boundaries (Sukumaran and Knowles 2017; MacGuigan et al. 2021). Lastly, integrating over gene trees by MCMC is computationally demanding and limits the number of loci that can be analysed efficiently (Leaché et al. 2014). The growing availability of genome-wide data has therefore driven demand for implementations that scale to large datasets (Fujita et al. 2012; Leaché et al. 2014; Douglas et al. 2022). Two leading platforms, BEAST2 (Bayesian evolutionary analysis by sampling trees; Bouckaert et al. 2019) and BPP (Bayesian Phylogenetics & Phylogeography; Yang and Rannala 2010, 2014; Flouri et al. 2020), operate within this framework and offer distinct strategies for species delimitation and hypothesis testing (Table 1).

**Table 1.** Overview of Multispecies Coalescent (MSC)-based species delimitation and species tree estimation methods implemented in the software platforms Bayesian Phylogenetics & Phylogeography (BPP) and Bayesian evolutionary analysis by sampling trees (BEAST2).

| Programme | BPP |  |  |  | BEAST2 |  |  |
| --- | --- | --- | --- | --- | --- | --- | --- |
| Method / Algorithm | A00 | A01 | A10 | A11 | BFD*<br>(SNAPPER) | SPEEDEMON<br>(SNAPPER) | SPEEDEMON<br>(StarBeast3) |
| Hypothesis testing | ✓ |  |  |  | ✓ |  |  |
| Species discovery (*de novo*) |  |  |  | ✓ |  | ✓ | ✓ |
| Species delimitation<br>(fixed tree) |  |  | ✓ |  |  |  |  |
| Parameter estimation ($\theta$, $\tau$) | ✓ | ✓ | ✓ | ✓ | ✓ | ✓ | ✓ |
| Species tree inference |  | ✓ |  | ✓ | ✓ | ✓ | ✓ |
| Data type | Sequence<br>alignments | Sequence<br>alignments | Sequence<br>alignments | Sequence<br>alignments | Biallelic<br>markers | Biallelic<br>markers | Sequence<br>alignments |
| Required *a priori* species<br>assignments | ✓ | ✓ | ✓ |  | ✓ |  |  |
| Clock model | Strict,<br>Relaxed | Strict,<br>Relaxed | Strict,<br>Relaxed | Strict,<br>Relaxed | Strict | Strict | Strict,<br>Relaxed |
| Gene flow models | ✓ |  |  |  |  |  |  |
| Calibrations (tip dating) | ✓ | ✓ | ✓ | ✓ |  |  | ✓ |
| Morphology (implementation traits) | ✓ |  |  |  |  |  | ✓ |
| Parallelisation | ✓ | ✓ | ✓ | ✓ |  |  | ✓ |

$\tau$ = divergence time; $\theta$ = effective population size; MSC-I, multispecies coalescent with introgression; MSC-M, multispecies coalescent with migration.

Bayes Factor Delimitation (BFD*) (Leaché et al. 2014) is a BEAST2 implementation designed for formal Bayes factor comparisons among competing species delimitation hypotheses from genomic data. It builds on SNAPP (Bryant et al. 2012) and its successor SNAPPER (Stoltz et al. 2021), which estimate species trees directly from biallelic markers such as single nucleotide polymorphisms (SNPs). SNAPP analytically integrates over gene tree topologies under the MSC using a pruning algorithm, whereas SNAPPER replaces this with a diffusion approximation of the Wright-Fisher model (Donnelly and Kurtz 1996; Stoltz et al. 2021), reducing computational demand while retaining the coalescent framework. Using these implementations, BFD* obtains marginal likelihood estimates by path sampling, enabling the hypothesis comparisons to species *a priori*, but BFD* does not depend on a specified topology, avoiding potential bias from the tree structure (Bryant et al. 2012; Leaché et al. 2014).

SPEEDEMON (SPEciEs DEliMitatiON, Douglas and Bouckaert 2022) is another BEAST2 implementation, offering a more heuristic approach to delimitation. Unlike BFD*, it can be run not only with *a priori* species assignments but also under ’unrestricted species discovery’, in which each individual starts as a putative species and the method explores possible configurations. SPEEDEMON is based on a birth-death collapse model, originally developed in DISSECT (Jones et al. 2015) and STACEY (Jones 2017), with an extension that allows the speciation rate to vary through time. It places a ’spike-and-slab’ prior on divergence times, governed by a user-defined threshold (ε): nodes above ε follow a standard birth-death tree prior (the ’slab’), while those below are collapsed and treated as conspecific (the ’spike’).

Similar to BFD*, SPEEDEMON is built on SNAPPER and can approximate species boundaries from biallelic data without estimating gene trees. More recent implementations extend it to multilocus sequence data and relaxed clock models through StarBeast3 (Douglas and Bouckaert 2022; Douglas et al. 2022); although StarBeast3 adds the computational cost of gene tree estimation, this is offset by optimised MCMC operators and parallelisation across threads (Douglas et al. 2022).

BPP is a Bayesian program that works exclusively with multilocus sequence data and integrates over gene trees by MCMC throughout. It implements four model-algorithms: A00 estimates population parameters, i.e., node heights (τ) and effective population size per branch (θ), under a fixed species tree (Rannala and Yang 2003; Flouri et al. 2020); A01 infers the species tree when species assignments are fixed (Rannala and Yang 2017); A10 performs species delimitation under a fixed guide tree (Yang and Rannala 2010; Rannala and Yang 2013); and A11 jointly infers species boundaries and tree topology (Yang and Rannala 2014). Like SPEEDEMON, A11 allows *de novo* species discovery, but instead of moving node heights across a collapse threshold it uses reversible-jump MCMC to switch between discrete species configurations. All BPP algorithms require a guide tree as a starting point, which A00 and A10 hold fixed and A11 is free to depart from. Under A00, BFdriver enables marginal likelihood estimation, allowing Bayes factor comparisons among competing hypotheses analogous to BFD*, but with marginal likelihoods obtained via Gauss-Legendre quadrature (Rannala and Yang 2017). A00 has also been extended to model gene flow, through the MSC-I model for episodic introgression and the MSC-M model for continuous migration (Flouri et al. 2020). Distinguishing these approaches by data type, inference strategy, treatment of species hypotheses, and underlying assumptions allows researchers to choose the method best suited to their study system.

The methods outlined above are increasingly applied to taxa with established morphological frameworks and broadly accepted species hypotheses, such as birds (Musher and Cracraft 2018), insects (Duran et al. 2024), amphibians (Scott et al. 2025), and reptiles (Leaché et al. 2009), but remain underused in marine invertebrates (Ramírez-Portilla and Quattrini 2023), where diagnostic features are often limited or highly plastic. For these organisms, key parameters such as effective population sizes, mutation rates, and divergence times are often unknown (Schuster et al. 2018; Ramírez-Portilla and Quattrini 2023), and they inhabit environments without clear physical barriers to gene flow, blurring patterns of genetic differentiation and species boundaries (Knowlton 1993; Palumbi 1994). Even where MSC-based methods have been applied to such groups, no study has systematically compared the major Bayesian implementations on a single empirical dataset to identify both their performance differences and the methodological factors driving them.

Sponges (Phylum Porifera) are a concrete example of a taxonomically complex marine invertebrate group. These sessile filter feeders are widely distributed across nearly all benthic aquatic ecosystems and fulfil numerous critical ecological roles (Bell 2008; De Goeij et al. 2013), but species distinction within the group remains notoriously difficult. Species identification still rests largely on macroscopic (e.g., colour, texture, habitus) and microscopic characters (e.g., the shape, type, and arrangement of skeletal elements; Payne et al. 2026), many of which are highly plastic and vary seasonally or locally with temperature, silicate concentration, and water flow (Maldonado et al. 1999; Morrow et al. 2013), leading to subjective and inconsistent classifications (Dohrmann et al. 2006; Erpenbeck et al. 2025). Mediterranean golf ball sponges (*Tethya* spp.), however, are a notable exception, with established morphological and ecological criteria for distinguishing species (Cardone et al. 2010; Corriero et al. 2015), making them a suitable proof-of-concept system for evaluating MSC-based delimitation in a group known for its complex systematics.

Three species of *Tethya* currently co-occur in the lagoon Mar Piccolo of Taranto, Italy (Ionian Sea, Central Mediterranean): *T. citrina*, *T. meloni*, and *T. aurantium* (Fig. 1). *T. citrina* was subsumed within *T. aurantium* for almost two centuries despite obvious morphological differences (Pallas 1766; Sarà and Melone 1965), until allozyme data confirmed that they form distinct genetic lineages (Sarà et al. 1989). The three differ in external morphology (body size, colour), skeletal characters (particularly the megasters), reproductive mode, and ecological niche, providing several independent lines of evidence for their distinction (Table S1; Corriero et al. 2015). *T. citrina* occurs in the mesolittoral zone under rocks in shallow water (∼1 m), where it is regularly exposed to air at low tide; *T. aurantium* is restricted to low-disturbance, low-light microhabitats such as small cavities in rock embankments, with a patchy distribution; and *T. meloni*, the most widespread of the three, colonises vertical artificial structures and retaining walls, forming dense aggregations of up to 100 individuals per m² (Cardone et al. 2010). *T. citrina* and *T. aurantium* frequently reproduce asexually by budding, a strategy rarely observed in *T. meloni* (Corriero et al. 2015). These ecological and reproductive differences are likely to limit gene flow among the three species, consistent with their long-recognised genetic distinctness (Sarà et al. 1989). This well-characterised system allows us to test whether MSC-based delimitation recovers species boundaries consistent with the established morphological and ecological framework. Using target capture enrichment of single-copy nuclear loci with a tethyid-specific bait set (see Erpenbeck et al. 2025), we assess the performance and congruence of the different approaches. A departure from the expected three species would indicate either that morphology-based taxonomy fails to capture the true boundaries, or that some implementations are sensitive to violations of model assumptions such as ongoing gene flow or population structure. We further evaluate the influence of topological constraints, data type, and clock model on the outcomes, and explore the potential role of introgression. Where the methods converge or conflict, we examine the underlying drivers and ask whether the resulting species hypotheses are robust and biologically meaningful.

**Figure 1.**
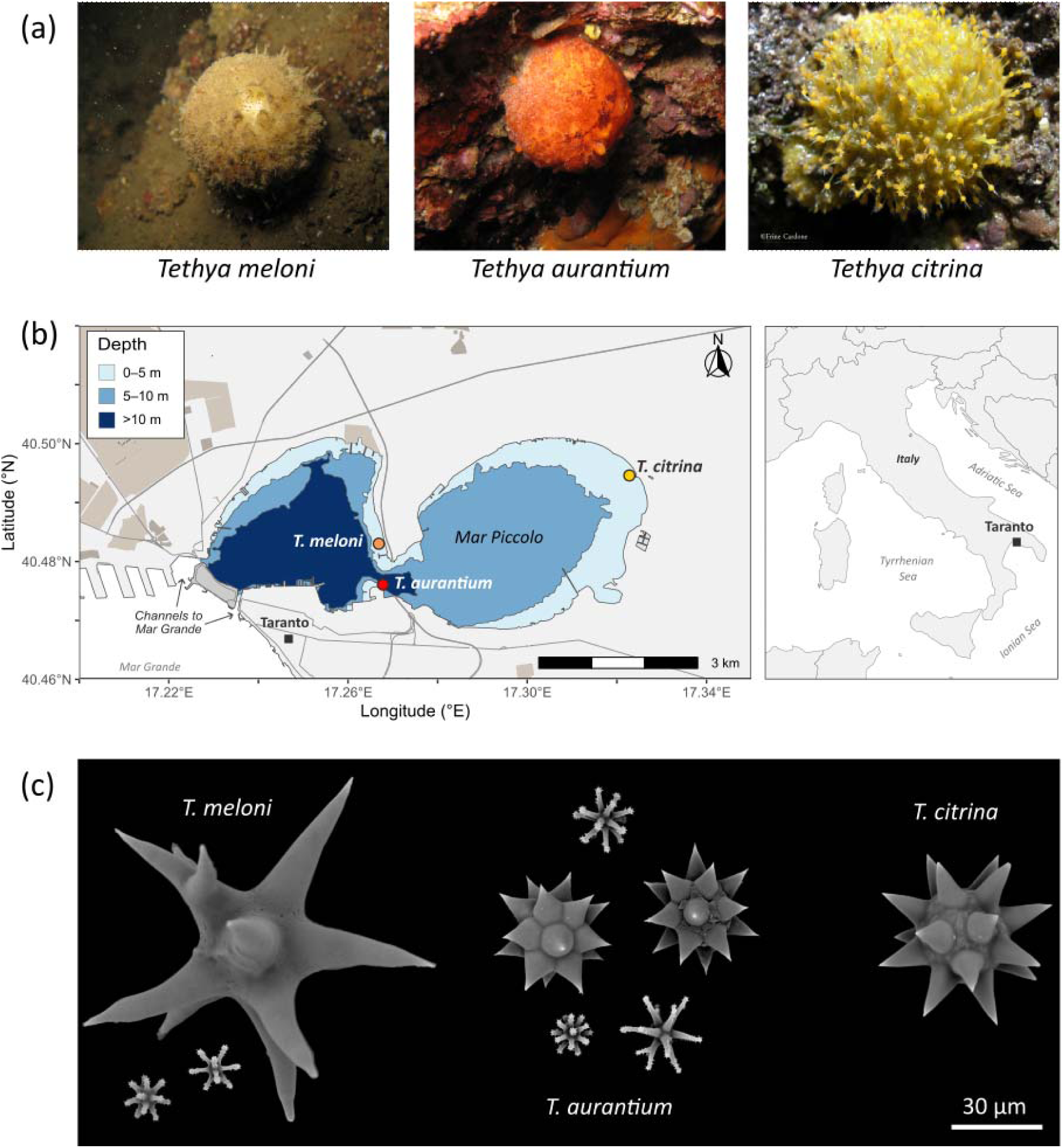
**(a)** *In situ* photographs of *Tethya meloni*, *T. aurantium*, and *T. citrina* (left to right). **(b)** Distribution of the three Mediterranean *Tethya* species in the shallow lagoon Mar Piccolo of Taranto, Italy (Ionian Sea, Central Mediterranean). **(c)** Scanning electron micrographs (SEM) of species-diagnostic megasters of *T. meloni* (left), *T. aurantium* (centre), and *T. citrina* (right). Scale bars is 30 µm. *In situ* and SEM photographs: © Frine Cardone.

## Material and Methods

### Sampling of Tethya Specimens

Specimens of the three Mediterranean *Tethya* species were collected in Mar Piccolo (40°30’07.17"N, 17°15’47.61"E), a semi-enclosed coastal lagoon at the northern end of the Gulf of Taranto, Italy (Ionian Sea, Central Mediterranean). In total, 44 specimens were sampled: 12 *T. aurantium*, 13 *T. meloni*, and 19 *T. citrina*. Two non-Mediterranean specimens were included as outgroup taxa: *T. wilhelma* (GW33333), from an aquarium population at Ludwig-Maximilians-Universität München, Germany (Wörheide et al. 2024), and *T. seychellensis* (GW41675), from the intertidal zone of Magoodhoo, Faafu Atoll, Maldives (collected and exported under permit (OTHR)30-D/INDIV/2017/379, Ministry of Fisheries and Agriculture, Maldives). *T. wilhelma* has affinities to Indo-Pacific *Tethya* (Sarà et al. 2001), and *T. seychellensis* is an Indo-Pacific species; as their taxonomic distinction is ambiguous (Erpenbeck et al. 2025), they were treated as a single outgroup. Small tissue samples were excised from each specimen and immediately fixed in absolute ethanol. Specimens were morphologically identified to species level following Corriero et al. (2015) (Table S1).

### Molecular Data Acquisition

Genomic DNA was extracted from the *Tethya* specimens with Qiagen’s Genomic Tip Kit following the manufacturer’s protocol. Low molecular weight fragments were removed by size selection (bead/extract ratio 1/3) with AMPure XP beads (Beckman Coulter, USA). DNA quality and quantity were assessed on a Nanodrop 1100 (Thermo Fisher, USA), and fragment size was visualised on 1.5% agarose gels. Double-indexed paired-end libraries were prepared with Illumina’s Nextera kit or Swift Bioscience’s 2S Library Kit, quantified on a Qubit 2.0 fluorometer with the dsDNA High Sensitivity Assay Kit (Invitrogen, USA), and checked on a Bioanalyzer 2100 with the High Sensitivity DNA Kit (Agilent, USA).

Target capture enrichment followed the myBaits protocol (Arbor Biosciences, USA; hybridisation temperature 62 °C), using a custom set of tethyid-specific baits targeting 7,755 single-copy nuclear genes (Erpenbeck et al. 2025). Four multiplexed equimolar libraries were pooled per hybridisation reaction. Post-hybridisation libraries were amplified with KAPA HiFi polymerase (Roche), checked on a Bioanalyzer 2100, and sequenced (150PE) on an Illumina MiniSeq.

### Bioinformatic Processing

Raw reads were quality-filtered and trimmed with *fastp* (settings: -q 28, -g -X -n 0; Chen et al. 2018) and assembled with metaSPAdes under default settings (Nurk et al. 2017). Bait sequences were mapped to each assembly with LASTZ (Harris 2007), allowing multiple mapping so that contigs matching more than one bait could be identified and excluded; only contigs uniquely mapped to a single bait per sample were retained. Loci were grouped by bait, aligned with MAFFT (Katoh et al. 2002), and trimmed with GBlocks 0.91b (Castresana 2000; Talavera and Castresana 2007), using default thresholds for conserved and flanking positions (50% and 85% of sequences, respectively), a minimum of 15 contiguous non-conserved positions, gaps allowed at all positions, and the default minimum block length of 10. Loci present in ≥ 95% of samples were concatenated into a supermatrix for ML inference in RAxML 8.2.12 (Stamatakis 2014).

### Variant Calling

Reads from each specimen were mapped to the targeted contigs of *T. wilhelma* (GW33333) with BWA-MEM (Li 2013) under default settings. The resulting BAM files were sorted with samtools (Li et al. 2009), duplicates were marked with MarkDuplicates in Picard (http://broadinstitute.github.io/picard), and reads were grouped per sample with Picard’s AddOrReplaceReadGroups. Haplotypes were called per sample with GATK v4.6.1 HaplotypeCaller, combined into a database with GenomicsDBImport, and jointly genotyped with GenotypeGVCFs (McKenna et al. 2010). The resulting VCF was filtered with VariantFiltration to remove SNPs meeting any of the standard GATK hard-filter thresholds (QD < 2.0, FS > 60.0, MQ < 40.0, MQRankSum < −12.5, ReadPosRankSum < −8.0). Further stringent filtering in bcftools (Li 2011) retained only biallelic SNPs passing depth and quality criteria (MIN(FORMAT/DP) >= 10; FILTER = "PASS"; N_ALT == 1; TYPE == "SNP"), leaving 2,066 SNPs. The VCF was converted to NEXUS format with *vcf2nex.pl* (https://www.beast2.org/snapp-faq/).

### MSC-Based Species Delimitation Analyses

#### BFD*

Species hypothesis testing was performed using BFD* (Leaché et al. 2014) implemented in BEAST2 v2.7.7 (Bouckaert et al. 2019) with SNAPPER v1.1.5 (Stoltz et al. 2021). Species assignments followed the ML phylogeny (Fig. 2), from which we defined six putative species groups: *T. citrina* divided into two clusters, *T. aurantium* divided into one divergent individual (GW1941) and the remaining specimens, *T. meloni* as a single group, and *T. wilhelma* and *T. seychellensis* combined as the outgroup. From these groups, five delimitation hypotheses were proposed: H3, a conservative lumping scenario merging *T. aurantium* and *T. meloni* into one species; H4, the three morphologically defined Mediterranean species (*T. citrina*, *T. meloni*, *T. aurantium*); H5.1, which additionally splits *T. citrina* into two clusters; H5.2, which instead splits *T. aurantium* into the divergent individual and the remaining specimens; and H6, treating all six groups as distinct. The outgroup was retained in all hypotheses.

**Figure 2.**
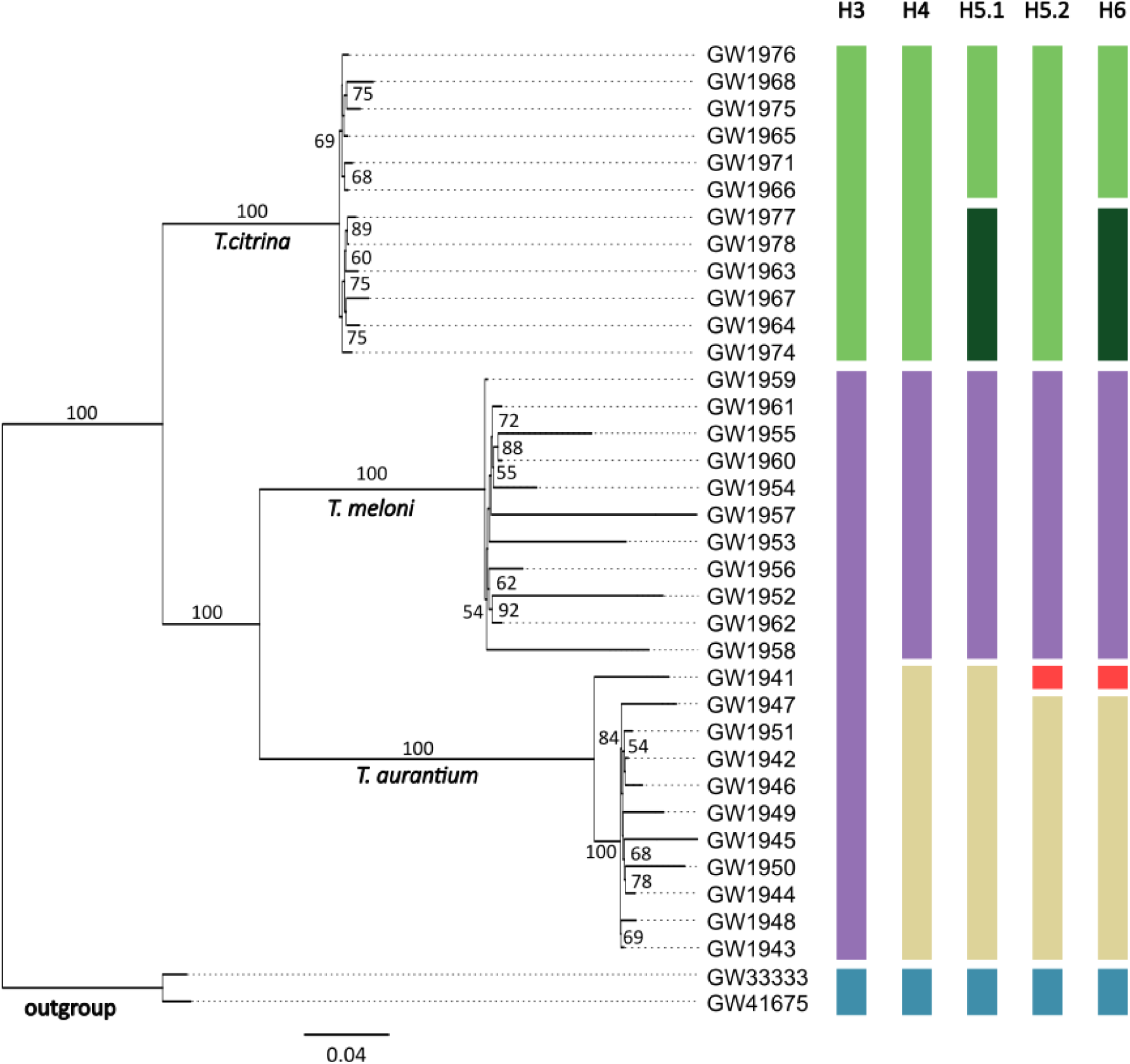
Maximum likelihood phylogeny (26,214 bp) of the 34 Mediterranean *Tethya* specimens with sufficient genomic data, based on 64 concatenated single-copy nuclear loci and rooted on the outgroup specimens *T. wilhelma* (GW33333) and *T. seychellensis* (GW41675). Scale is substitutions/site; node values are bootstrap probabilities (BP, in %), shown only for BP ≥ 50. Coloured bars denote the species delimitation hypotheses (H3, H4, H5.1, H5.2, H6).

Path sampling used 100 steps, a chain length of 5,000,000 generations per step, a pre-burnin of 1,000,000 generations, and a sampling burn-in of 50%. The Alpha parameter was set to 0.3 (default; shape of the Beta-distributed inverse temperature schedule) and a strict clock was applied with default settings. Given the lack of reliable calibration data and the general uncertainty about divergence times in demosponges (Schuster et al. 2018), we specified broad, weakly informative gamma priors (SNAPPER coalescent rate ∼ Gamma(α = 2, β = 1); Yule birth rate ∼ Gamma(α = 2, β = 2)). Marginal likelihood estimates (MLEs) were obtained with PathSampleAnalyser, and Bayes factors are reported as 2×ln(BF), calculated as 2 × (MLE[ − MLE[) on the log scale. On this scale, values of 0–2 indicate no support, 2–6 moderate, 6–10 high, and >10 very high support (Kass and Raftery 1995).

#### SPEEDEMON

SPEEDEMON v1.1.0 (Douglas and Bouckaert 2022) was applied in BEAST2 using both SNAPPER and StarBeast3 v1.2.1 (Douglas et al. 2022) in BEAST2. Two species assignment strategies were tested: an *a priori* assignment (H6; Fig. 2) and an unrestricted species discovery (USD) approach (Douglas and Bouckaert 2022). Both implementations used a Yule Skyline Collapse tree prior with 4 epochs (equal lengths, linked means; default settings), collapse weight ∼ Beta(α = 3, β = 1), and birth rate ∼ LogNormal(M = 1.0, S = 1.25, meanInRealSpace = true). The collapse threshold ε is expressed in expected substitutions per site, and the appropriate value differs between implementations. SNP-based tree heights in SNAPPER are larger than the sequence-based estimates in StarBeast3 and therefore require higher ε. As the developers advise against fixing ε at its default (Douglas and Bouckaert 2022), a range of values was tested for each method.

For SNAPPER, the coalescent rate prior was set to Gamma(α = 2, β = 2) under a strict clock. We tested ε = 0.075, 0.100, 0.125, and 0.150 under the a priori assignment (H6), and repeated the USD analysis at the best-performing ε. For StarBeast3, we tested ε = 0.005, 0.001, 0.0005, and 0.0001 under H6 with a strict clock, comparing linked and unlinked site models to assess the sensitivity of delimitation to site model configuration. Under linked models, all loci share the GTR+Gamma parameters; under unlinked models, each locus has independent parameters. At the best-performing ε, four further analyses were run: H6 and USD, each under linked and unlinked site models with a strict clock, plus H6 with linked site models under a relaxed clock.

For both methods, MCMC chains were run for 100,000,000 generations with a pre-burnin of 1,000,000. Posterior distributions of species assignments were summarised with the ClusterTreeSetAnalyser tool using a 10% burn-in.

#### BPP Parameter Estimation, Species Tree Estimation and Species Delimitation

Parameter estimation, species tree estimation, and species delimitation were performed using BPP v4.8.4 (Yang and Rannala 2010, 2014; Flouri et al. 2020). We applied four BPP algorithms: A00 (speciesdelimitation = 0, speciestree = 0) for parameter estimation under a fixed species tree, A01 (speciesdelimitation = 0, speciestree = 1) for species tree estimation under fixed species assignments, A10 (speciesdelimitation = 1, speciestree = 0) for species delimitation under a fixed guide tree, and A11 (speciesdelimitation = 1, speciestree = 1) for joint species delimitation and tree inference using rjMCMC. Specimens were assigned to six putative species groups (H6) based on the ML inference (Fig. 2): *T. aurantium* was split into two groups (A, n = 1; B, n = 10), *T. meloni* formed a single group (C, n = 11), *T. citrina* was split into two groups (D, n = 6; E, n = 6), and the two outgroup specimens formed one group (F, n = 2). The fixed guide tree was specified as (F,((D,E),(C,(A,B)))).

For A01, an outgroup constraint was applied (F: *T. wilhelma* and *T. seychellensis*), and for A01, A10, and A11 the speciesmodelprior was set to 1 (uniform prior on rooted trees). All analyses used 64 loci under the GTR substitution model with among-locus rate variation (alphaprior = 1 1 4; gamma prior with shape α = 1, mean = 1, and 4 categories). For the initial analyses, inverse-gamma (IG) priors were placed on the mutation-scaled effective population size (θ) and divergence time (τ): θ ∼ IG(3, 0.01) and τ ∼ IG(3, 0.001), with prior means of 0.005 and 0.0005, respectively. The shape parameter α = 3 followed the BPP manual (Yang 2015), with β values calibrated to the low substitution rates expected in sponges (Wörheide et al. 2012; Schuster et al. 2018). All four algorithms were run under both a strict clock (default) and a relaxed clock with independent rates among loci (clock = 2; μ = 10.0, ν = 100.0, σ² = 5.0). To assess prior sensitivity, the strict-clock analyses were repeated under two further configurations: a broad prior (θ ∼ IG(3, 0.1), mean 0.05; τ ∼ IG(3, 0.01), mean 0.005) and a narrow prior (θ ∼ IG(3, 0.002), mean 0.001; τ ∼ IG(3, 0.0003), mean 0.00015).

Each MCMC was run for 1,000,000 iterations with a 100,000-generation burn-in, sampling every 10 generations, on 32 threads. Unrestricted species discovery, in which each individual is treated as a putative species, was not implemented in BPP, as preliminary benchmarking indicated that exploring the full species configuration space under A11 was computationally prohibitive, with an expected runtime of at least six months.

#### BFdriver

Bayes factor comparisons among species delimitation hypotheses were performed using BFdriver in BPP (Rannala and Yang 2017). We tested the same five hypotheses (H3, H4, H5.1, H5.2, H6; see Fig. 2). All settings were identical to the A00 analysis under the strict clock and initial priors (θ ∼ IG(α = 3, β = 0.01); τ ∼ IG(α = 3, β = 0.001)). Because marginal likelihood estimation requires multiple quadrature points per hypothesis and is computationally costly, however, each MCMC was run for 100,000 samples after a burn-in of 10,000 generations, sampling every 10 generations on 16 threads. Log marginal likelihoods were obtained by summing the weighted expected log-likelihoods across all quadrature points, and Bayes factors were calculated as the ratio of marginal likelihoods and interpreted following Kass and Raftery (1995).

#### Introgression Models MSC-I

Introgression among Mediterranean *Tethya* was assessed using the MSC-I method (A00) in BPP, following the parameterisation of Flouri et al. (2020). Four models were tested: (1) a ghost lineage model (Fig. 3a), in which introgression coincided with the parental divergence (τ-parent = yes, yes); (2) an incipient speciation model (Fig. 3b), in which introgression occurred after the parental divergence but the recipient lineage diverged at the time of introgression (τ-parent = no, yes); (3) a horizontal gene flow model (Fig. 3c), in which both the introgression event and the recipient divergence occurred independently of the parental node ages (τ-parent = no, no); and (4) a bidirectional gene flow model (Fig. 3d), in which gene flow occurred in both directions between two sister lineages, with initial introgression probabilities φ = 0.1 and φ = 0.2.

**Figure 3.**
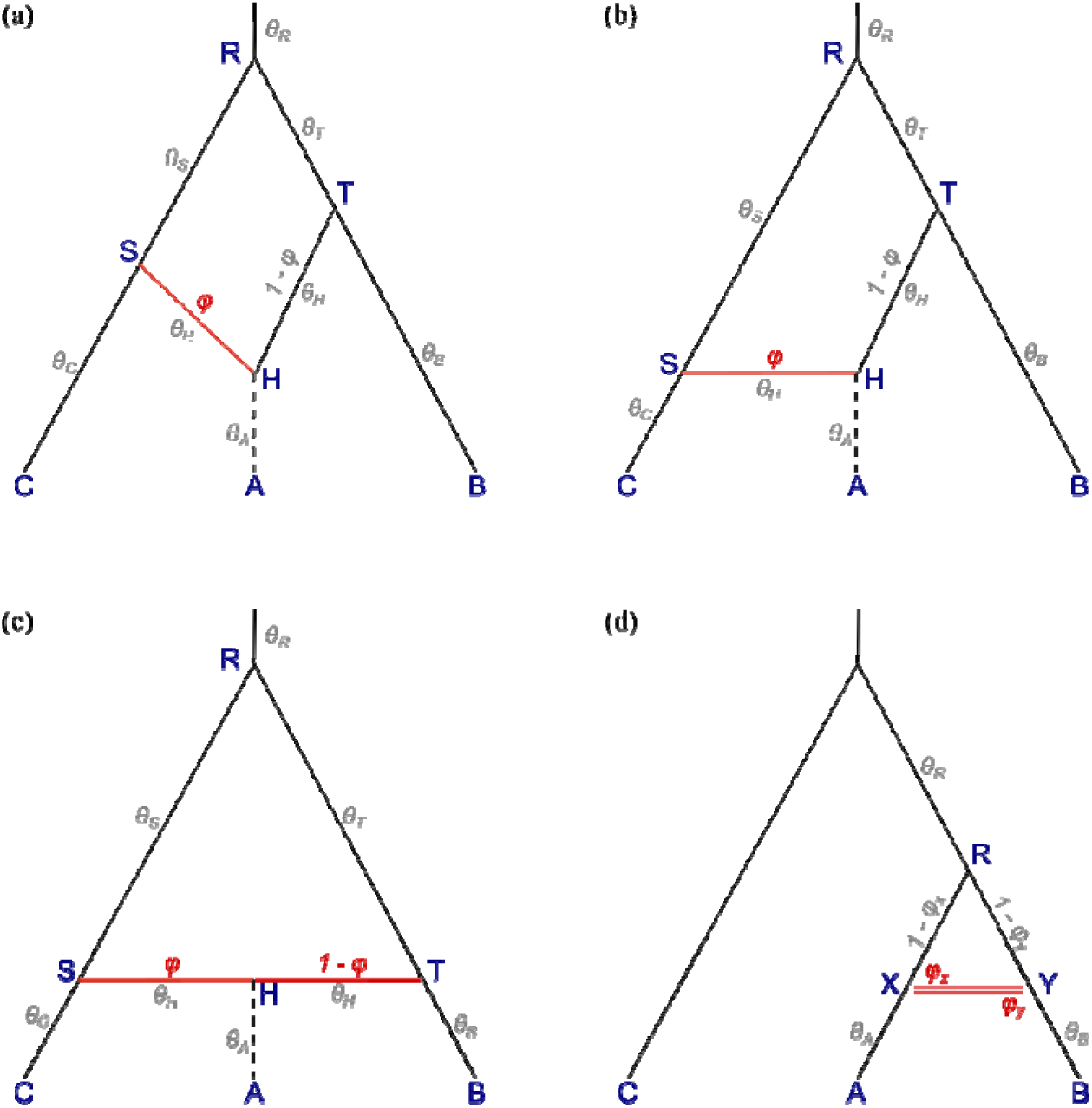
The four introgression models tested with the MSC-I method in BPP, adapted from Flouri et al. (2020). Tips A, B, and C represent extant species; internal nodes R, S, T, and H represent ancestral populations or hybridization nodes. Branch labels (θ) indicate population size parameters for each branch. Red lines indicate introgression events, with φ denoting the introgression probability (i.e., the proportion of genes contributed by the donor lineage) and 1 – φ the contribution of the parental lineage. (**a**) Ghost lineage model, where the introgression coincides with parental divergence. (**b**) Incipient speciation model, where introgression occurs after parental divergence. (**c**) Horizontal gene flow model, where introgression occurs independently of parental divergence times. (**d**) Bidirectional gene flow model, with reciprocal introgression between sister lineages A and B (φ_x_ and φ_y_).

Under the four-species hypothesis (H4), all four models were applied, with tips A, B, and C corresponding to *T. aurantium*, *T. meloni*, and *T. citrina*, respectively. Under the six-species hypothesis (H6), only the ghost lineage and incipient speciation models were tested, with tip A corresponding to *T. aurantium* sp. 1 (GW1941), B to *T. aurantium* sp. 2, and C to *T. meloni*. In all analyses, φ was assigned a flat Beta(1, 1) prior, and all other settings were identical to the A00 analysis under the strict clock and initial priors. Marginal likelihoods and Bayes factors were estimated using BFdriver as described above.

All MSC-based analyses were run in duplicate to assess consistency between independent runs. Convergence was assessed using Tracer v1.7.2 (Rambaut et al. 2018) by examining trace plots and ESS values (> 200), supplemented by inspection of acceptance rates and operator diagnostics. All data, control files, and scripts used in this study are available at https://github.com/PalMuc/SpongeMSCSD.

## Results

### Genomic Dataset Assembly

Raw reads were retrieved for 38 of the 44 collected specimens, with six yielding insufficient DNA for library preparation. After quality filtering and *de novo* assembly, four further specimens were excluded for low read coverage, leaving a final dataset of 34 Mediterranean and two outgroup specimens (Table S2). Target capture recovered between 5,398 and 14,519 single-copy nuclear loci per sample (mean: 8,741 ± 1,858 SD), of which 64 were present in ≥ 95% of samples and used for downstream analyses. Variant calling and filtering retrieved 2,066 SNPs.

### Species Hypothesis Testing

Both BFD* and BFdriver identified H6 as the best-supported hypothesis among the five candidates (Fig. 2; Table 2), with very strong support under both methods (2×lnBF > 10). Both ranked H5.2 above H5.1, indicating stronger support for the *T. aurantium* split than for the *T. citrina* split, while H3, which lumps *T. aurantium* and *T. meloni*, received the lowest support. Replicates produced consistent MLEs across methods, and all analyses converged (ESS > 200).

**Table 2.** Bayes factor testing of five candidate species delimitation hypotheses under two methods: BFD* (SNAPPER; BEAST2) with SNP data and marginal likelihood estimates (MLEs) from path sampling, and BFdriver (BPP) with multilocus sequence data and MLEs from Gauss-Legendre quadrature. Bayes factors (2×ln BF = 2 × (MLEbest − MLEalternative); Kass and Raftery 1995) are 0 for the best-supported hypothesis, H6, and positive for each alternative, with values > 10 indicating strong support for H6 over that alternative.

| Hypothesis | #Species | BFD* |  |  | BFdriver |  |  |
| --- | --- | --- | --- | --- | --- | --- | --- |
|  |  | MLE | BF | Rank | MLE | BF | Rank |
| H3 | 3 | -53,277.82 | 36,189.37 | 5 | -135,547.16 | 2,046.34 | 5 |
| H4 | 4 | -35,926.20 | 1,486.11 | 4 | -134,687.08 | 326.18 | 4 |
| H5.1 | 5 | -35,665.83 | 965.39 | 3 | -134,606.90 | 165.82 | 3 |
| H5.2 | 5 | -35,446.15 | 526.05 | 2 | -134,577.28 | 106.58 | 2 |
| H6 | 6 | -35,183.13 | 0 | 1 | -134,523.99 | 0 | 1 |

### De Novo Species Delimitation and Discovery

#### SPEEDEMON-SNAPPER

All analyses reached convergence (ESS > 200) and produced consistent results across replicates. However, the inferred species delimitation varied with the collapse threshold ε, shifting from H5.2 (PP ≥ 0.999) at lower ε (ε = 0.075, 0.100) to H4 (PP ≥ 0.998) at higher ε (ε = 0.150; Fig. 4), with a transition at ε = 0.125 where both hypotheses received moderate but incomplete support (H4: PP = 0.78; H5.2: PP = 0.22). Cluster support for H5.2 and H4 was distributed across multiple species tree topologies (Table S3a). In the ultrametric species tree, the outgroup was nested as sister to *T. meloni*, with *T. citrina* sister to that clade and *T. aurantium* the earliest-branching lineage (Fig. 4a). This topology was best supported at ε = 0.075, 0.100, and 0.150 (PP = 0.83–0.85) and least at ε = 0.125 (PP = 0.65), where five topologies were needed to cover the 95% credible set.

**Figure 4.**
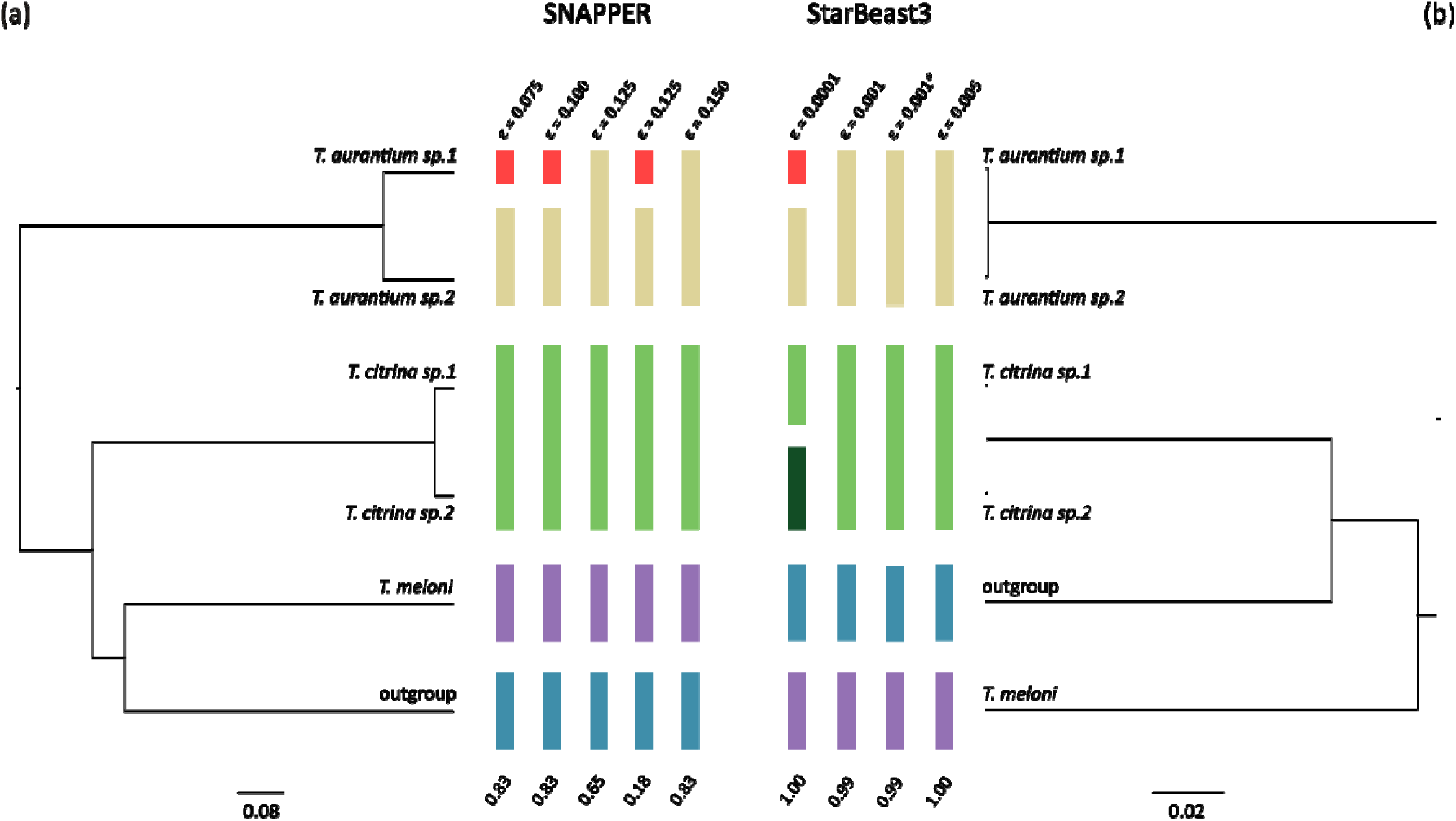
SPEEDEMON species delimitation for Mediterranean *Tethya* under a strict clock model, using *a priori* species assignments (six-species hypothesis H6; see Fig. 2). Coloured bars show the species clusters recovered at each collapse threshold (ε); bars of the same colour denote merged lineages, and values below each column give the posterior probability (PP) of the corresponding delimitation and species-tree hypothesis. Displayed species trees are ultrametric maximum clade credibility (MCC) trees (median node heights; scale bars in expected substitutions (mutation-scaled) per site, see Table 3 for Tree.Height values). (**a**) SPEEDEMON-SNAPPER, tree shown for ε = 0.125. (**b**) SPEEDEMON-StarBeast3, tree shown for ε = 0.001. Site-model parameters were linked across loci; the analysis marked with an asterisk (*) was additionally run with unlinked parameters.

**Table 3.** SPEEDEMON parameter estimates under varying collapse thresholds (ε). (**a**) SNAPPER with SNP data; (**b**) StarBeast3 with sequence data under a strict clock; (**c**) StarBeast3 under a relaxed clock. H6 = *a priori* species assignment; USD = unrestricted species discovery. Linked/Unlinked indicates whether site model parameters were shared across loci. Posterior, lnL, and prior are natural-log values; estimates are posterior means. Asterisks (*) mark analyses that did not converge.

| | | $\epsilon$ | Posterior | lnL | Prior | Cluster Count | Collapse Weight | Tree Height |
| --- | --- | --- | --- | --- | --- | --- | --- | --- |
| **(a) SNAPPER** |  |  |  |  |  |  |  |  |
| **H6** |  | 0.075 | -35133.63 | -35084.63 | -49.01 | 5.000 | 0.445 | 0.759 |
|  |  | 0.100 | -35133.67 | -35084.39 | -49.28 | 4.999 | 0.444 | 0.759 |
|  |  | 0.125 | -35132.31 | -35084.84 | -47.47 | 4.216 | 0.531 | 0.756 |
|  |  | 0.150 | -35131.67 | -35084.32 | -47.35 | 4.002 | 0.556 | 0.759 |
| **USD** |  | 0.125 | -33037.82 | -32836.81 | -201.01 | 6.506 | 0.833 | 0.766 |
| **(b) StarBeast3 (Strict Clock)** |  |  |  |  |  |  |  |  |
| **H6** | Linked | 0.0001 | -1.24E+05 | -1.31E+05 | -38.30 | 6.000 | 0.332 | 0.130 |
|  |  | 0.0005* | -1.23E+05 | -1.31E+05 | -33.66 | 4.113 | 0.545 | 0.118 |
|  |  | 0.001 | -1.24E+05 | -1.31E+05 | -29.42 | 4.006 | 0.557 | 0.128 |
|  |  | 0.005 | -1.24E+05 | -1.31E+05 | -30.97 | 4.000 | 0.555 | 0.132 |
|  | Unlinked | 0.001 | -1.24E+05 | -1.30E+05 | -826.46 | 4.000 | 0.554 | 0.141 |
| **USD** | Linked | 0.001* | -1.22E+05 | -1.31E+05 | 152.392 | 6.663 | 0.829 | 0.123 |
|  | Unlinked | 0.001* | -1.22E+05 | -1.30E+05 | -649.842 | 7.854 | 0.798 | 0.129 |
| **(c) StarBeast3 (Relaxed Clock)** |  |  |  |  |  |  |  |  |
| **H6** | Linked | 0.001* | -1.23E+05 | -1.30E+05 | -41.193 | 4.962 | 0.499 | 0.090 |

### SPEEDEMON-StarBeast3

All analyses converged (ESS > 200) and produced consistent results across replicates, except for ε = 0.0005. The species delimitation varied with ε, shifting from H6 (PP = 1.0) at ε = 0.0001 to H4 at ε = 0.001 (PP = 0.994) and ε = 0.005 (PP = 1.0), with negligible support for H5.2 at any threshold (PP ≤ 0.003). The species relationships themselves, however, were consistent across all analyses, with *T. citrina* grouped with the outgroup and *T. meloni* with *T. aurantium* (Table S3b), but the position of the root within this topology was poorly resolved. At ε = 0.001, the best-supported rooting placed *T. aurantium* as the earliest-branching lineage (Fig. 4b), but with low support (PP = 0.42), and two alternative rootings received comparable support (PP = 0.30 and 0.27; Table S3b). This contrasts with SNAPPER, which recovered a stable root (*T. aurantium* earliest-branching) but variable relationships among the remaining lineages. Unlinked site models at ε = 0.001 recovered the same delimitation and relationships (H4, PP > 0.99; Table S3c).

### SPEEDEMON: Relaxed Clock Analyses

Relaxed clock analyses failed to converge (ESS < 200), owing to poor mixing of individual gene tree heights (particularly in R2); other parameters, such as per-locus clock rates, birth rates, and species tree height, mixed adequately (ESS > 200). The recovered species relationships were consistent across replicates and matched the strict-clock analyses, but the posterior probabilities of the delimitation hypotheses fluctuated within and between replicates (Fig. S1). The coefficient of variation of species tree branch rates was 1.0–1.2 across replicates (ESS > 200), indicating rate heterogeneity among branches.

### SPEEDEMON

#### Unrestricted Species Discovery

Unrestricted species discovery failed to converge under both SPEEDEMON implementations. SNAPPER (ε = 0.125), which does not support BEAGLE acceleration, completed only ∼30% of the target 100M states after more than three months of computation, and convergence diagnostics improved only marginally with continued sampling. Global parameters mixed adequately (e.g., Tree.height ESS = 1,959–2,389; BirthRates ESS > 19,000; clusterCount ESS = 319–987), but likelihood-related parameters remained far below the 200 threshold (posterior ESS = 23–39; likelihood ESS = 29–33), as did most per-locus snapperCoalescentRate and theta parameters (ESS < 50).

StarBeast3, which supports BEAGLE acceleration and multithreading, completed the full 100M-state chain within 3.5–5.5 days on 32 threads across all tested configurations (ε = 0.001, linked and unlinked site models). However, all analyses failed to converge and gave inconsistent posterior distributions between replicates. ESS values for core parameters (posterior, likelihood, species coalescent, clusterCount, SPEEDEMONYuleSkylineCollapse) remained below 60 in the best-performing runs and below 10 in the worst. Linear extrapolation indicates that reaching ESS > 200 for the slowest-mixing parameters would require a 4- to 50-fold increase in chain length, projecting a minimum runtime of seven months under optimistic scaling. Across both implementations, unrestricted species discovery did not result in converged solutions within a feasible runtime. Non-converged cluster support values are shown in Figure S2 for completeness but cannot be interpreted as reliable delimitations.

### BPP Parameter Estimation, Species Delimitation, and Species Tree Inference

#### A00 Parameter Estimation

A00 analyses converged under both clock models (ESS > 200) with consistent results across replicates. Strict-clock divergence times (τ) ranged from the root (τ_root = 0.070) to the shallowest splits within *T. aurantium* (τ_AB = 0.00032) and *T. citrina* (τ_DE = 0.00071; Table 4a). Mutation-scaled effective population sizes (θ) differed among lineages, with the estimate for *T. meloni* (θ_C = 0.034) roughly ten times larger than those for the *T. aurantium* and *T. citrina* sublineages (θ_B, θ_D, θ_E; Table 4a). The ancestral population size for the MRCA of *T. meloni* and *T. aurantium* (θ_CAB = 1.79) was disproportionately high relative to all other nodes.

**Table 4.** Population size (θ) and divergence time (τ) estimates from BPP A00 under an *a priori* six-species assignment (H6). Guide tree: (F,((D,E),(C,(A,B)))). Values represent posterior means with 95% HPD (Highest Posterior Density) intervals. (**a**) Initial prior setting under strict and relaxed clock models; (**b**) broad prior setting under strict clock; (**c**) narrow prior setting under strict clock. MRCA = most recent common ancestor.

| (a) $\theta \sim \text{IG}(\alpha = 3, \beta = 0.01)$; $\tau \sim \text{IG}(\alpha = 3, \beta = 0.001)$ | | | | | | | | | |
| --- | --- | --- | --- | --- | --- | --- | --- | --- | --- |
|  |  | Strict Clock |  |  |  | Relaxed Clock |  |  |  |
| Node | Label | $\theta$ | $\theta$ (95% HPD) | $\tau$ | $\tau$ (95% HPD) | $\theta$ | $\theta$ (95% HPD) | $\tau$ | $\tau$ (95% HPD) |
| A | *T. aurantium* sp.1 | - | - | - | - | - | - | - | - |
| B | *T. aurantium* sp.2 | 0.0035 | 0.0016–0.0055 | - | - | 0.0056 | 0.0037–0.0077 | - | - |
| C | *T. meloni* | 0.0336 | 0.0285–0.0389 | - | - | 0.2594 | 0.1977–0.3219 | - | - |
| D | *T. citrina* sp.1 | 0.0037 | 0.0025–0.0049 | - | - | 0.0040 | 0.0027–0.0054 | - | - |
| E | *T. citrina* sp.2 | 0.0072 | 0.0047–0.0099 | - | - | 0.0066 | 0.0044–0.0090 | - | - |
| F | Outgroup | 0.0234 | 0.0175–0.0296 | - | - | 0.0193 | 0.0139–0.0252 | - | - |
| A, B | MRCA *T. aurantium* | 0.0208 | 0.0164–0.0256 | 0.00032 | 0.00010–0.00057 | 0.0633 | 0.0507–0.07765 | 0.0011 | 0.00063–0.00162 |
| D, E | MRCA *T. citrina* | 0.0116 | 0.0095–0.0138 | 0.00071 | 0.00047–0.00094 | 0.0255 | 0.0208–0.0306 | 0.00081 | 0.00055–0.00109 |
| C, A, B | MRCA *T. meloni* + *T. aurantium* | 1.7904 | 0.4002–4.3212 | 0.00593 | 0.00499–0.00689 | 0.0089 | 0.0009–0.0264 | 0.08983 | 0.07878–0.10120 |
| D, E, C, A, B | MRCA ingroup | 0.8818 | 0.7256–1.0492 | 0.01166 | 0.00909–0.01443 | 0.0093 | 0.0009–0.0265 | 0.08985 | 0.07879–0.10121 |
| Root | MRCA all | 0.4162 | 0.3610–0.4716 | 0.06974 | 0.06177–0.07754 | 0.3590 | 0.2721–0.4516 | 0.08987 | 0.07880–0.10123 |

**(b)** $\theta \sim \text{IG}(\alpha = 3, \beta = 0.1)$ ; $\tau \sim \text{IG}(\alpha = 3, \beta = 0.01)$
| Node | Label | Strict Clock |  |  |  |
| --- | --- | --- | --- | --- | --- |
| | | $\Theta$ | $\Theta$ (95% HPD) | $\tau$ | $\tau$ (95% HPD) |
| A | *T. aurantium* sp.1 | - | - | - | - |
| B | *T. aurantium* sp.2 | 0.0067 | 0.0048–0.0088 | - | - |
| C | *T. meloni* | 0.0340 | 0.0289–0.0393 | - | - |
| D | *T. citrina* sp.1 | 0.0056 | 0.0042–0.0071 | - | - |
| E | *T. citrina* sp.2 | 0.0102 | 0.0070–0.0137 | - | - |
| F | Outgroup | 0.0250 | 0.0189–0.0316 | - | - |
| A, B | MRCA *T. aurantium* | 0.0206 | 0.0160–0.0255 | 0.00068 | 0.00039–0.00098 |
| D, E | MRCA *T. citrina* | 0.0114 | 0.0094–0.0134 | 0.00091 | 0.00068–0.00114 |
| C, A, B | MRCA *T. meloni* + *T. aurantium* | 1.8270 | 0.3956–4.1674 | 0.00595 | 0.00501–0.00689 |
| D, E, C, A, B | MRCA ingroup | 0.8850 | 0.7298–1.0533 | 0.01166 | 0.00909–0.01443 |
| Root | MRCA all | 0.4172 | 0.3624–0.4734 | 0.06988 | 0.06180–0.07765 |

**(c)** $\theta \sim \text{IG}(\alpha = 3, \beta = 0.002)$ ; $\tau \sim \text{IG}(\alpha = 3, \beta = 0.0003)$
| Node | Label | Strict Clock |  |  |  |
| --- | --- | --- | --- | --- | --- |
| | | $\Theta$ | $\Theta$ (95% HPD) | $\tau$ | $\tau$ (95% HPD) |
| A | *T. aurantium* sp.1 | - | - | - | - |
| B | *T. aurantium* sp.2 | 0.0021 | 0.0003–0.0043 | - | - |
| C | *T. meloni* | 0.0345 | 0.0284–0.0417 | - | - |
| D | *T. citrina* sp.1 | 0.0035 | 0.0023–0.0047 | - | - |
| E | *T. citrina* sp.2 | 0.0069 | 0.0044–0.0096 | - | - |
| F | Outgroup | 0.0232 | 0.0174–0.0294 | - | - |
| A, B | MRCA *T. aurantium* | 0.0215 | 0.0165–0.0274 | 0.00018 | 0.00002–0.00042 |
| D, E | MRCA *T. citrina* | 0.0114 | 0.0084–0.0139 | 0.00067 | 0.00043–0.00092 |
| C, A, B | MRCA *T. meloni* + *T. aurantium* | 1.5407 | 0.0004–3.8281 | 0.00616 | 0.00500–0.00789 |
| D, E, C, A, B | MRCA ingroup | 0.8966 | 0.7225–1.0870 | 0.01114 | 0.00692–0.01419 |
| Root | MRCA all | 0.4164 | 0.3614–0.4722 | 0.06949 | 0.06118–0.07738 |

The relaxed clock indicated rate heterogeneity among loci (mean locus rate variance ν[ = 0.615; 95% HPD: 0.511–0.721). Here, the divergence times at the three deepest nodes collapsed to comparable values (τ = 0.090), θ_CAB decreased drastically from 1.79 to 0.009, and lnL was higher than under the strict clock (−129,540 vs. −130,031). Strict-clock estimates were otherwise robust to alternative prior configurations (Table 4b, c), with point estimates stable across deeper nodes and the largest differences at the shallowest nodes, where 95% HPD intervals broadened under the narrow prior and the lower bound of τ_AB extended to near-zero. ESS values remained > 200 under both configurations.

### A01 Species Tree Estimation

All strict-clock analyses gave stable root parameter estimates (θ_root = 0.41–0.42; τ_root = 0.07). Under the initial priors, both replicates converged on the same maximum a posteriori (MAP) topology (Fig. 5a; PP = 0.74; lnL = −130,030), with the leading alternative placing *T. meloni* with *T. citrina* (PP = 0.24; Table S4). The broad prior failed to converge, each replicate recovering a different topology. The narrow prior returned the same MAP topology but with reduced support (PP = 0.40) and greater weight on alternatives, including *T. aurantium* with *T. citrina* (PP = 0.40) and *T. meloni* with *T. citrina* (PP = 0.20; Table S4). The relaxed clock instead favoured a topology placing *T. meloni* closer to the outgroup (Fig. 5b; PP = 1.0), with comparable θ_root (0.35) but higher τ_root (0.10) and lnL (−129,476).

**Figure 5.**
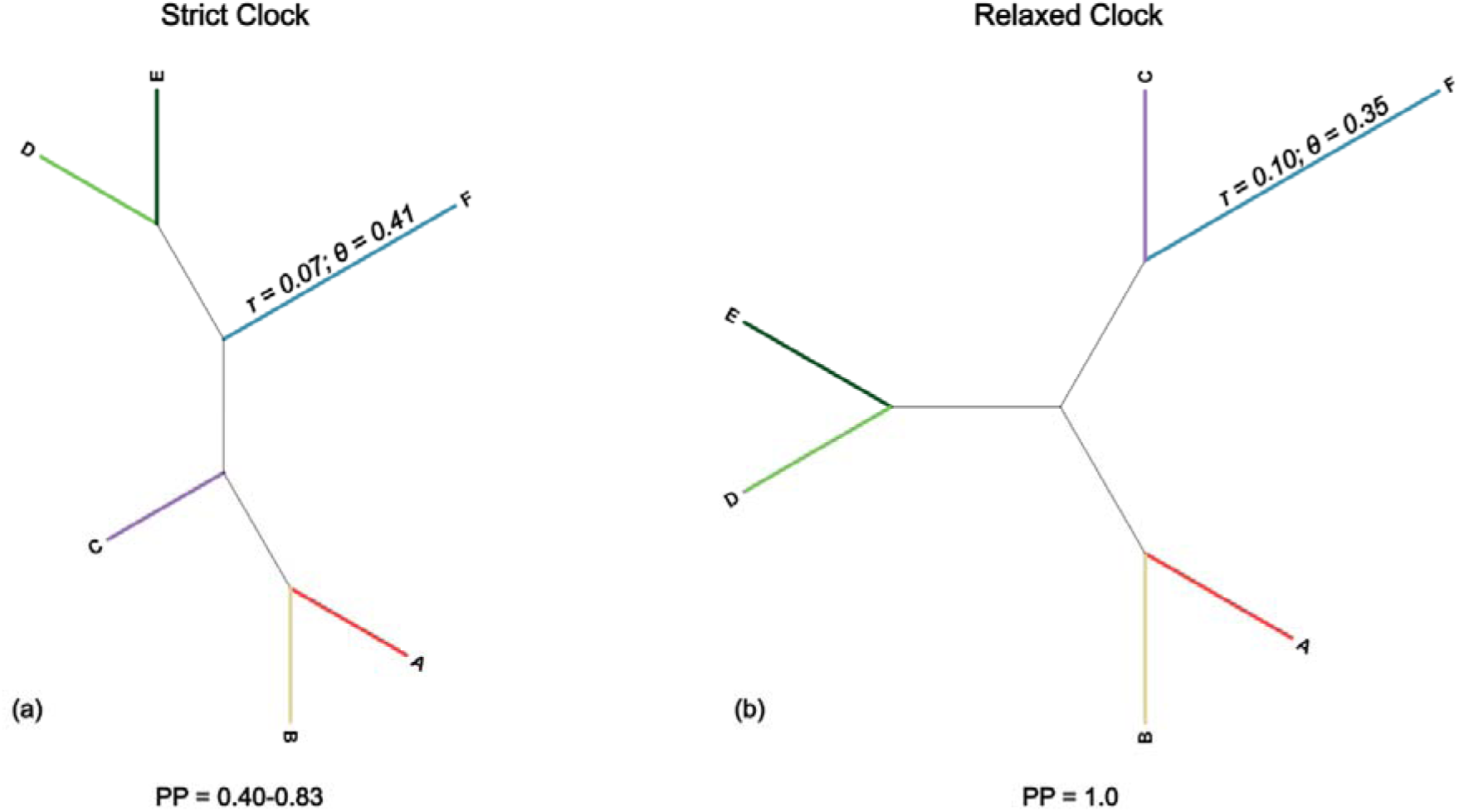
Species tree topologies inferred by BPP A01 under (**a**) a strict clock (initial and narrow prior settings) and (**b**) a relaxed clock (initial prior settings) model. Species assignments were fixed under the six-species hypothesis (H6), with the outgroup (F) constrained; the root estimates of effective population size (θ) and divergence time (τ) are shown. Tip labels: A = *T. aurantium* sp.1; B = *T. aurantium* sp.2; C = *T. meloni*; D = *T. citrina* sp.1; E = *T. citrina* sp.2; F = outgroup. Posterior probabilities (PP) of the maximum a posteriori topology are shown below each tree (see Table S4 for all sampled). Branch lengths are not to scale.

### A10 Species Delimitation using a Fixed Guide Tree

The A10 analyses recovered H6 with PP = 1.0 across all prior configurations and both clock models, converging within replicates (ESS > 200) and giving consistent results across replicates. Under the strict clock, parameter estimates were similar across all prior configurations (θ_root = 0.42, τ_root = 0.07), with lnL = −130,035, −130,072, and −130,028 for the initial, broad (θ ∼ IG(3, 0.1); τ ∼ IG(3, 0.01)), and narrow (θ ∼ IG(3, 0.002); τ ∼ IG(3, 0.0003)) priors, respectively. Under the relaxed clock, estimates were congruent but with a lower θ_root (0.36), a higher τ_root (0.09), and a higher lnL (−129,540 vs. −130,035 for the strict clock under the initial prior).

### A11 Joint Species Delimitation and Species Tree Estimation

The A11 analyses recovered H6 with PP = 1.0 across all prior configurations under the strict clock, despite a prior disadvantage for the six-species model (induced marginal prior of 0.130 for six species and 0.174–0.217 for three to five species; (Flouri et al. 2018)). Under the initial and broad priors, both replicates recovered a consistent species-tree topology, with *T. aurantium* and *T. meloni* as sisters and *T. citrina* grouped with the outgroup (F; θ_root = 0.37, τ_root = 0.09; PP = 1.0); here the outgroup was nested within the ingroup rather than forming the earliest-branching lineage. Under the narrow prior, the analyses failed to converge: R2 recovered a similar topology (PP = 1.0; lnL = −130,056), whereas R1 recovered an alternative placing *T. aurantium* sister to a *T. meloni* and *T. citrina* clade (PP = 0.9997; lnL = −130,115).

The A11 analyses under the relaxed clock also failed to converge. In R1, the chain entered an unrealistic parameter space, with τ_root drifting from 0.30 to 2.09 over the run and the log probability of the gene trees under the MSC (log-PG, as reported by BPP) exceeding 10¹²; this replicate recovered a topology with the outgroup sister to *T. meloni*. R2 was more stable (τ_root = 0.086; lnL = −129,540) and recovered a MAP tree with the outgroup as the earliest-branching lineage. H6 was recovered in both replicates (PP = 1.0), but the parameter and topology estimates from these analyses cannot be reliably interpreted.

### Introgression and Gene Flow

The MSC-I analyses under H4 produced inconsistent outcomes across models and replicates (Table 5). The ghost lineage model (Fig. 3a) gave comparable introgression probabilities (φ) between replicates (R1: φ = 0.829; R2: φ = 0.880), but τ and θ at the introgression node varied, and R2 mixed far more poorly than R1 (ESS = 258 vs. 108,881). The incipient speciation (Fig. 3b) and horizontal gene flow (Fig. 3c) models converged, but in the incipient speciation model τ_root and τ at the introgression node (τ_intro) collapsed to comparable values (0.075 and 0.071, respectively), leaving no temporal separation between them. The horizontal gene flow model estimated φ = 0.993 (95% HPD: 0.980–1.000). The bidirectional gene flow model (Fig. 3d) failed to converge, with R1 supporting asymmetrical gene flow from *T. aurantium* to *T. meloni* (φ = 0.306; 95% HPD: 0.265–0.347) and R2 estimating this parameter close to zero (φ = 0.009; 95% HPD: 0.000–0.022).

**Table 5.**
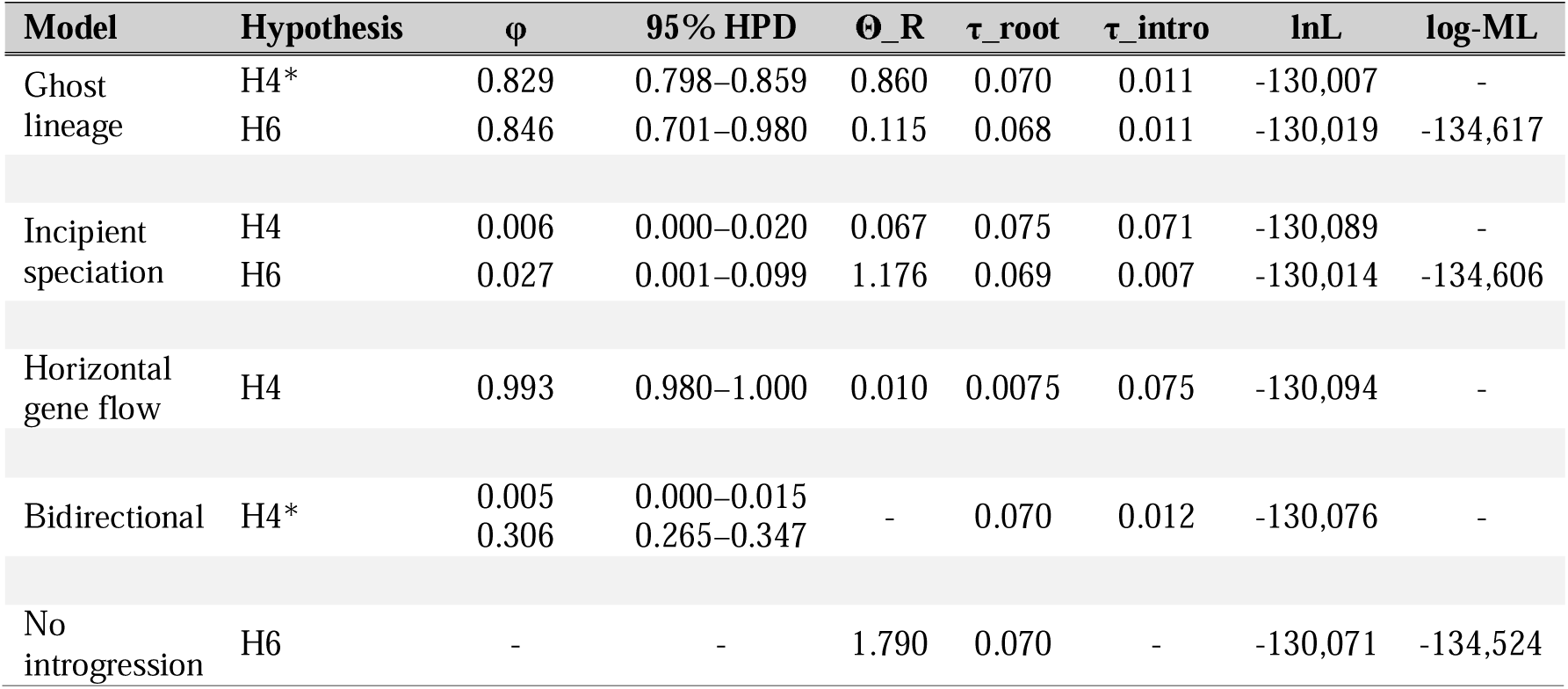
BPP introgression model (MSC-I) results under the four-species (H4) and six-species (H6) hypotheses. φ = introgression probability (posterior mean with 95% HPD interval); θ_R = ancestral population size at the introgression node; τ_root and τ_intro = divergence times at the root and introgression node, respectively; lnL = log-likelihood; log-ML = log marginal likelihood estimated via BFdriver. Analyses marked with an asterisk (*) showed inconsistent parameter estimates between replicates (R1 estimates are displayed).

Under H6, both the ghost lineage and incipient speciation models converged across replicates. The ghost lineage model estimated a high introgression probability (φ = 0.846; 95% HPD: 0.714–0.980), reducing the ancestral population size at the introgression node from 1.790 (A00) to 0.115, whereas the incipient speciation model estimated low introgression (φ = 0.027; 95% HPD: 0.001–0.099) but retained a high ancestral population size (θ_R = 1.176). Both achieved higher MCMC log-likelihoods than H6 without introgression (lnL = −130,014 and −130,019 vs. −130,071 for A00), yet MLE via BFdriver favoured H6 without introgression (log-ML = −134,524) over both the incipient speciation (−134,606; 2×lnBF = 164) and ghost lineage models (−134,617; 2×lnBF = 186; Table 5).

A full overview of all model output, MCMC trace files, and control files the MSC-based analyses reported are available in the GitHub repository (https://github.com/PalMuc/SpongeMSCSD).

## Discussion

Species delimitation methods based on the multispecies coalescent are increasingly applied across the tree of life, but how they behave in taxonomically difficult, non-model groups such as sponges remains poorly understood. Mediterranean *Tethya* offer a rare opportunity to address this, as a group whose species-level boundaries are independently established on morphological and ecological grounds, against which the output of each method can be assessed. Applied to this system, all MSC-based methods consistently recovered *T. aurantium*, *T. citrina*, and *T. meloni* as distinct lineages, regardless of analytical framework, clock model, prior configuration, or collapse threshold. Genetic divergence between them is sufficiently deep to corroborate the established morphological taxonomy (Sarà and Melone 1965; Corriero et al. 2015). At the sublineage level, however, the methods diverged. The constrained, hypothesis-testing approaches converged on the same six-species hypothesis (H6) across different data types and model assumptions, whereas the more exploratory approaches were less stable and, in several configurations, limited by the signal in the loci or by computational cost. Here, we discuss why the methods differ in their output, how much confidence each result warrants, and which configurations remain workable in a system of this kind, providing a reference for studies facing comparable systematic complexity.

### Hypothesis-Testing Approaches Converge on H6

All three hypothesis-testing approaches applied here (BFD*, BFdriver, and BPP A10) recovered H6 as the best-supported delimitation, despite differing in inference strategy, guide-tree dependence, and data type within their shared constrained framework. A10 uses rjMCMC on multilocus sequences, BFdriver Gauss-Legendre quadrature on the same data, and BFD* path sampling on biallelic SNPs via SNAPPER. Furthermore, A10 and BFdriver rely on a guide tree, whereas BFD* does not. Differences like these can produce divergent outcomes, and BPP and BFD* have previously been shown not to always recover congruent delimitations when applied to the same system (Ferrer Obiol et al. 2023). Their agreement on H6 here, together with its consistent recovery across both clock models in A10, indicates that the 64 loci and the biallelic SNPs each carry sufficient signal to detect genetic differentiation, and that this outcome is robust to data type, guide-tree choice, and inference strategy.

### Prior Sensitivity, Guide Tree and Clock Models in BPP

The recovery of H6 extended to the full set of BPP analyses, regardless of prior configuration or clock model. BPP is known to be sensitive to prior settings, particularly the prior on effective population sizes (Leaché and Fujita 2010; Yang 2015). In our analyses, parameter estimates were largely consistent across prior configurations at deeper nodes, with only modest shifts at shallow nodes (Table 4a–c). At these shallow nodes, limited genetic differentiation reduces the information available to distinguish intraspecific from interspecific variation; thus, the prior exerts proportionally more influence on the estimates (Yang and Rannala 2014).

We did notice that species tree estimation under A01 was more prior-sensitive than the delimitation algorithms (A10, A11), which may be the result of the outgroup constraint we imposed. Because BPP infers a rooted species tree, we placed F (*T. wilhelma* and *T. seychellensis*) at the root of A01 using the available biogeographic information. However, the unconstrained A11 analyses suggest that this rooting conflicts with the data. Namely, under the strict clock, A11 recovered a tree in which the outgroup grouped with *T. citrina* instead of forming the earliest-branching lineage, and this topology was consistent under both the initial and broad priors. This agrees with earlier work showing that deep relationships within *Tethya* remain poorly resolved (Sorokin et al. 2019; Santodomingo et al. 2024), and indicates that biogeographic isolation or distinctness can be a poor guide for rooting a species tree.

Despite these prior-driven differences in parameter estimates and topology, the delimitation outcome itself was robust. H6 received the highest posterior probability across all converged analyses, including under A11 despite a disadvantage for the six-species model under the uniform rooted-tree prior. The 64 loci therefore carry sufficient signal to overcome this disadvantage and to support the six putative lineages regardless of prior and clock model, matching the expectation that weak signals across loci accumulate under the MSC to reveal lineage structure undetectable from a single locus (Degnan and Rosenberg 2009; Jiao et al. 2021). The A11 relaxed-clock analyses, on the other hand, failed to converge, with τ_root drifting, log-PG inflating in R1, and topologies differing between replicates. H6 was still nominally recovered, but these estimates cannot be reliably interpreted. Hence, with more complex, heuristic algorithms, sensitivity to the prior and to model assumptions increases, and this effect is amplified under the relaxed clock.

### Rate Heterogeneity Among Lineages

The ancestral population size of the MRCA of *T. meloni* and *T. aurantium* (θ_CAB) was disproportionately high across all prior configurations (Table 4a). Under the MSC, such inflation can arise either from rate heterogeneity among lineages not accommodated by the clock model (Burgess and Yang 2008; Flouri et al. 2022) or from unmodelled gene flow between the descendant lineages (Tiley et al. 2023), since in both cases the excess substitutions accumulating in the ancestral population are absorbed into θ. We tested both possibilities, first by comparing strict and relaxed clock models and subsequently through MSC-I models.

The relaxed clock confirmed rate heterogeneity among lineages, with a coefficient of variation in branch rates well above zero (νC = 0.615; 95% HPD: 0.511–0.721) and a higher log-likelihood than the strict clock. Most notably, θ_CAB dropped from 1.79 to 0.009, while the three deepest divergence times converged on near-identical values (τ = 0.090, with overlapping HPD intervals; Table 4a). The inflated θ_CAB under the strict clock therefore largely absorbed unmodelled among-lineage rate variation, and is less likely to be indicative of a large ancestral population. For parameter estimation under a fixed tree (A00), the relaxed clock captures this rate variation better and removes the inflation, but it is also more parameter-rich. Under A11, where the species tree is inferred jointly, the weak signal in the loci at the deep nodes leaves too little information to estimate the topology and the per-branch rates together, causing convergence to fail.

This τ-collapse indicates that the deepest nodes approach a star-like configuration, in which short internal branches offer few opportunities for lineage sorting and hierarchical relationships cannot be reliably resolved under the MSC fo this system. Consistent with this, the inferred resolution at the deep nodes depended on the clock model. Under the relaxed clock, A01 placed *T. meloni* closer to the outgroup than under the strict clock (Fig. 5), and A11 under the relaxed clock failed to converge. Both patterns are expected under a short-internal-branch regime, where weak deep-node signal leaves the topology poorly determined. Although this complicates parameter estimation, it is biologically plausible for *Tethya* and supports an early, rapid radiation of the genus, originally proposed from patterns of morphological and biogeographic diversity (Sarà and Sarà 2004) and reflected in the weak biogeographic concordance of the main *Tethya* clades in molecular phylogenies (Santodomingo et al. 2024).

### Testing Ancestral Gene Flow (MSC-I)

To evaluate whether unmodelled gene flow could contribute to the inflated θ_CAB, we tested MSC-I models under both H4 and H6 (Tiley et al. 2023). However, none produced biologically interpretable results. Under H4, the bidirectional and ghost lineage models failed to converge across replicates, with τ and θ at the introgression node differing between runs (Table 5). The incipient speciation and horizontal gene flow models did converge, but their τ estimates at the root and introgression node collapsed to comparable values, leaving no temporal separation between parental divergence and introgression. The horizontal gene flow model further estimated φ = 0.993, implying that nearly all genetic information in *T. aurantium* derived from the donor lineage, which is more likely the result of a model misfit than a reliable signal.

Under H6, both the ghost lineage and incipient speciation models converged across replicates (Table 5), but their parameter estimates remain unrealistic. The ghost lineage model estimated a high introgression probability (φ = 0.846; 95% HPD: 0.714–0.980) and reduced θ_CAB from 1.790 to 0.115. This value is still an order of magnitude higher than the 0.009 recovered under the relaxed clock, thus only part of the inflated signal shifted to the introgression parameter. The estimated φ would moreover imply that ∼85% of the genetic material in *T. aurantium* sp.1 derived from an unsampled donor lineage, requiring either a recent massive introgression event or a donor representing the majority genomic background, neither of which is readily compatible with the genetic structure observed within the lagoon. Although the incipient speciation model estimated low introgression (φ = 0.027), it retained the inflated ancestral population size (θ_CAB = 1.176), leaving the inflation unresolved by introgression. BFdriver additionally favoured H6 without introgression over both MSC-I models, providing independent support for this conclusion.

Rate heterogeneity among lineages offers a more realistic explanation for the inflated θ_CAB than introgression, although introgression cannot be fully ruled out. Several limitations temper our ability to distinguish the two. The 64 loci may carry insufficient signal to estimate φ reliably, as hundreds to thousands of loci are normally recommended for robust estimation (Flouri et al. 2020). Furthermore, we acknowledge that the sampling under H6 is suboptimal, with n = 1 for *T. aurantium* sp.1 precluding within-lineage coalescent estimation of θ. Lastly, sampling was restricted to Mar Piccolo, therefore a donor lineage outside the lagoon cannot be ruled-out, particularly as closely related *Tethya* lineages are not necessarily geographically proximate (Santodomingo et al. 2024). And despite their apparently limited larval dispersal, this mismatch between dispersal potential and genetic structure is common in sponges and other marine invertebrates (Palumbi 1994; Maldonado 2006). Broader geographic and within-lineage sampling, together with more loci, could clarify whether rate heterogeneity is the dominant driver or whether introgression is also a contributing factor, though such analyses will likely face considerable computational bottlenecks at the moment.

### SPEEDEMON and the Choice of Collapse Thresholds

Among the methods tested, SPEEDEMON is attractive by design, exploring species configurations without *a priori* assignments and within a relatively short runtime, which makes it an apparently unbiased approach well suited to groups whose morphological characters are scarce or plastic. In our analyses, however, the outcome varied with ε, with low thresholds favouring more delimited species (H6 under StarBeast3, H5.2 under SNAPPER) and higher thresholds recognising only the three Mediterranean species (H4; Table 3, Fig. 4). This sensitivity contrasts with Douglas and Bouckaert (2022), who found ε to be robust across several orders of magnitude, although their analyses focused on vertebrates (geckos and primates). Hence, the difference may be system-dependent. In Mediterranean *Tethya* the splits of interest lie close to the collapse boundary, causing small shifts in ε tipping these nodes between the ’spike’ and the ’slab’ of the collapse prior. Varying the threshold thus reveals where each node falls relative to the boundary, but offers little insight into the underlying species boundaries in our system.

Although its developers characterise ε as a computational approximation to zero with no inherent biological significance (Douglas and Bouckaert 2022), applying it meaningfully would require an external benchmark for species formation, such as an undisputed reference species established independently at the relevant divergence depth. The threshold could then be calibrated for the focal group by setting it to a fraction of the smallest divergence between species (e.g., Voisin et al. 2025). For the *Tethya* sublineages, it remains unknown whether such a benchmark exists, as the sublineages may as well reflect population structure instead of species boundaries. The three Mediterranean species are firmly established but sit deeper than the splits at issue. In addition, sponges, like many marine invertebrates, lack a workable operational criterion such as reproductive isolation under the biological species concept (DeBiasse and Hellberg 2015; Shaffer et al. 2021). Although the hypothesis-testing approaches consistently recovered H6, using this as a reference is not a solution. Calibrating ε against the delimitations recovered by the other methods would only be circular, tuning SPEEDEMON towards an outcome it is meant to test independently. The choice of ε is thus arbitrary here, and exposes a gap in our knowledge of the study system.

### Data Type Shapes Species Tree Topology

Under a strict clock and *a priori* species assignments, SPEEDEMON-SNAPPER and SPEEDEMON-StarBeast3 recovered different topologies (Fig. 4), each resolving a different part of the tree. StarBeast3 recovered the same lineage relationships as the ML phylogeny (*T. citrina* with the outgroup, *T. meloni* with *T. aurantium*) but left the root poorly resolved, whereas SNAPPER recovered a stable root (*T. aurantium* earliest-branching) but relationships among the remaining lineages that did not match the ML topology. As both analyses used the same clock model and species assignments, this difference is most likely the result of how each implementation handles the input data. StarBeast3 jointly estimates a gene tree for each locus and constrains these within the species tree (Douglas et al. 2022), whereas SNAPPER analytically integrates over all possible gene trees using an allele-frequency diffusion model (Stoltz et al. 2021). Because StarBeast3 uses full sequences while SNAPPER relies on variable sites alone, and full sequences improve both branch-length and topological accuracy (Leaché et al. 2015), the SNP-based analysis in SPEEDEMON is placed at a disadvantage.

### Model Complexity and Computational Limits

The SPEEDEMON results discussed so far were obtained under the simplest configuration, combining a strict clock with *a priori* species assignments. Given the rate heterogeneity among lineages, a relaxed clock would be more appropriate for Mediterranean *Tethya* than a strict clock, and unrestricted species discovery (USD) would relax the *a priori* constraints; both, however, proved difficult to apply. Even the simplest relaxed-clock configuration in StarBeast3 (H6 with linked loci) showed convergence problems and inconsistent posterior distributions, although the estimated coefficient of variation in branching rates (∼1.0–1.2) indicated rate heterogeneity across lineages, consistent with the BPP results and supporting the use of a relaxed clock. Expanding the parameter space further impeded reliable inference, a pattern also seen in BPP A11 under the relaxed clock. However, SPEEDEMON analyses with independent per-locus parameters and those using USD likewise failed to converge under a strict clock (Table 3); thus, the problem is not confined to the clock model. Across the more complex models, the prior increased while log-likelihoods remained stable or decreased slightly, indicating that the posterior was increasingly driven by the prior and that model complexity exceeded the inferential capacity of the available loci.

USD removes the circularity of testing delimitations nested within a pre-specified hypothesis, but it also proved computationally intractable. Under SNAPPER, it traversed only ∼30% of the chain in over three months, with a projected runtime approaching a year, which is impractical for a method intended to be computationally efficient. StarBeast3 was faster but did not converge under a strict clock, with adequate convergence projected to require at least seven months, and a relaxed clock would only compound the problems already seen under H6. USD was not attempted in BPP A11, as preliminary benchmarking showed that exploring the full species configuration space was equally too intensive. The unbiased configurations we would have preferred, SPEEDEMON USD and BPP A11, could therefore not be reliably applied to our dataset.

### Toward MSC-Based Species Hypotheses to Inform Taxonomy

To our knowledge, this is the first systematic comparison of the major Bayesian MSC-based delimitation methods on a single empirical dataset, in a system long regarded as taxonomically challenging. The constrained, hypothesis-testing approaches recovered a congruent, robust and reproducible six-species delimitation. The heuristic and threshold-based approaches proved less stable. Knowing how and where the methods diverge is useful in practice, since it identifies constrained hypothesis-testing approaches as the most dependable in systems such as ours and shows where the others lose resolution. Model-based approaches yield quantifiable and replicable hypotheses that can be refined as more data become available.

This offers a more objective basis for defining species than the morphological interpretation of a single individual, occasionally supported by barcoding. Arguably, these methods are most needed in taxonomically complex groups whose morphology offers even fewer diagnostic characters than *Tethya*, such as Haplosclerida (Van der Sprong et al. 2025) and many other marine invertebrates with hard to define, plastic or reduced morphologies (Knowlton 1993; Ramírez-Portilla and Quattrini 2023).

Within the three recognised Mediterranean *Tethya* species, our results reveal further genetic structure. Its consistent recovery across methods makes a methodological artefact unlikely. Whether it represents species or populations remains to be resolved. MSC-based methods detect lineage structure but do not adjudicate species status (Leaché et al. 2019), and they can over-interpret clustering when population structure is present (Sukumaran and Knowles 2017; MacGuigan et al. 2021).

However, the observed substructure is biologically plausible here. The three *Tethya* species occupy patchy microhabitats within Mar Piccolo, and two of them reproduce frequently by asexual budding (Cardone et al. 2010; Corriero et al. 2015). Both factors are expected to reduce gene flow and promote differentiation (Wright 1943; Palumbi 1994). The resulting MSC-based delimitation can serve as a reference point and quantitative hypothesis against which other independent lines of evidence can be weighed to confirm or reject species status. By demonstrating how each method performs in a system challenged by its complex systematics, our study offers a reference for comparable systems and helps taxonomists make a better-informed choice of methodological approach. In doing so, it brings genome-scale insight to groups whose systematics have long resisted resolution.

## Author Contributions

JvdS, SV, DE, and GW were involved in the DFG-funded doctoral project. SV, DE, and GW designed the study and supervised the research. JvdS was primarily responsible for conducting the study. Supervision included assistance with data analyses, interpretation and manuscript preparation. FC contributed to the collection of the specimens and morphological identification of the specimens. SSC performed the laboratory work with support from JvdS and FD. FD designed the *Tethya* baits. JvdS, SV, SH led the implementation of the model-based species delimitation analyses. JvdS drafted the manuscript with substantial input from SV and SH. All authors read and approved the final version of the manuscript.

## Acknowledgements

We thank Gabriele Büttner and Nora Dotzler for their support in the lab, and René Neumaier for his assistance with the HPC infrastructure and for helping us run our analyses as efficiently as possible on the cluster. We further thank Wenjie Zhu and Luiza Guimarães Fabreti for their input on the computational aspects of this work, and Oliver Voigt for sharing his expertise on the study system. We would like to thank the staff of the MaRHE Center and the people of Magoodhoo (Faafu Atoll, Maldives) for support during field work. The Ministry of Fisheries and Agriculture, Maldives, is acknowledged for granting permission to carry out research in the Maldives. We are grateful to the editors and reviewers for their time and constructive feedback, which helped significantly to improve this manuscript.

## Data Availability Statement

Raw data are deposited in the European Nucleotide Archive (ENA) and accessible under the project PRJEB85726. All the model-based species delimitation analyses, workflows and input files can be found on our GitHub repository https://github.com/PalMuc/SpongeMSCSD, and archived at Zenodo: https://doi.org/10.5281/zenodo.20998531.

