## Supplementary Tables and Figures for "How Robust are Multispecies Coalescent Species Delimitations in Taxonomically Complex Systems? A Genomic Assessment Using Mediterranean *Tethya* Sponges"

**Table S1.** Comparative morphological and ecological traits of the three Mediterranean *Tethya* species (*T. aurantium, T. citrina, T. meloni*). Values are based on published data (primarily Corriero et al. 2015) and integrated with qualitative observations where quantitative measurements are not consistently available.

| **Character** | ***Tethya aurantium*** | ***Tethya citrina*** | ***Tethya meloni*** |
| --- | --- | --- | --- |
| Colour | Orange-red | Yellow | Cream / pale yellow |
| Body size | 1–4.5 cm | 0.5–2.5 cm | up to 8 cm |
| Cortex thickness | 1.5–3.5 mm; mean 2.6 mm | 1.0–1.5 mm; mean 1.2 mm | 3–5 mm; mean 4 mm |
| Cortex/body ratio | Intermediate | Low | High (up to ~1/3) |
| Consistency | Firm | Soft | Firm |
| Surface morphology | Papillate | Smooth / conulose | Flattened tubercles |
| Megaster size (µm) | 16–94; mean 54 (overlapping) | 18–65; mean 43 (overlapping) | 66–119; mean 90 (distinctly larger) |
| Megaster R/C ratio | ~0.5–0.6 | ~0.8–1.1 | ~1–1.8 |
| Micraster organisation | Structured (cortex *vs* choanosome) | Unstructured | Unstructured |
| Micraster types | Two categories (cortex vs choanosome) | Mixed types (oxyaster, chiaster, tylaster) | Highly heterogeneous (multiple types, variable size) |
| Asexual reproduction | Frequent (up to ~55 buds/ind.) | Very frequent (up to 250 buds/ind.) | Rare (2/89 individuals observed) |
| Habitat | Protected cavities | Intertidal / under rocks | Sheltered artificial structures / dense aggregations |

**Table S2**. Metadata and summary information of the *Tethya* specimens used in this study.

| **ID** | **Species** | **Total # Raw Reads** | **Total # Trimmed Reads** | **# Baits targeting SCG** | **# Contigs** | **BioProject** | **Reference** |
| --- | --- | --- | --- | --- | --- | --- | --- |
| GW1941 | *T. aurantium* | 212,506 | 201,776 | 6641 | 10144 | PRJEB85726 | This study |
| GW1942 | *T. aurantium* | 799,480 | 766,662 | 10003 | 16018 | PRJEB85726 | This study |
| GW1943 | *T. aurantium* | 487,688 | 467,090 | 9203 | 12277 | PRJEB85726 | This study |
| GW1944 | *T. aurantium* | 514,566 | 488,540 | 8128 | 15469 | PRJEB85726 | This study |
| GW1945 | *T. aurantium* | 763,052 | 719,876 | 7788 | 14907 | PRJEB85726 | This study |
| GW1946 | *T. aurantium* | 563,844 | 537,642 | 9074 | 13179 | PRJEB85726 | This study |
| GW1947 | *T. aurantium* | 357,388 | 332,566 | 7452 | 13916 | PRJEB85726 | This study |
| GW1948 | *T. aurantium* | 635,994 | 597,840 | 9194 | 22650 | PRJEB85726 | This study |
| GW1949 | *T. aurantium* | 349,738 | 330,138 | 7355 | 19219 | PRJEB85726 | This study |
| GW1950 | *T. aurantium* | 395,232 | 368,494 | 7702 | 19475 | PRJEB85726 | This study |
| GW1951 | *T. aurantium* | 552,776 | 530,282 | 8496 | 12729 | PRJEB85726 | This study |
| GW1952 | *T. meloni* | 1,218,426 | 1,173,432 | 5514 | 14143 | PRJEB85726 | This study |
| GW1953 | *T. meloni* | 615,160 | 585,156 | 6950 | 8864 | PRJEB85726 | This study |
| GW1954 | *T. meloni* | 740,328 | 705,574 | 7926 | 10334 | PRJEB85726 | This study |
| GW1955 | *T. meloni* | 484,846 | 463,392 | 6043 | 14404 | PRJEB85726 | This study |
| GW1956 | *T. meloni* | 694,812 | 665,100 | 7682 | 12380 | PRJEB85726 | This study |
| GW1957 | *T. meloni* | 644,864 | 616,244 | 5398 | 14879 | PRJEB85726 | This study |
| GW1958 | *T. meloni* | 761,588 | 726,634 | 6046 | 17545 | PRJEB85726 | This study |
| GW1959 | *T. meloni* | 592,092 | 567,826 | 8078 | 11492 | PRJEB85726 | This study |
| GW1960 | *T. meloni* | 590,432 | 558,668 | 8827 | 18743 | PRJEB85726 | This study |
| GW1961 | *T. meloni* | 894,724 | 894,636 | 8876 | 24314 | PRJEB85726 | This study |
| GW1962 | *T. meloni* | 882,588 | 849,824 | 8921 | 50185 | PRJEB85726 | This study |
| GW1963 | *T. citrina* | 573,370 | 546,166 | 9003 | 28629 | PRJEB85726 | This study |
| GW1964 | *T. citrina* | 902,608 | 861,052 | 9493 | 17642 | PRJEB85726 | This study |
| GW1965 | *T. citrina* | 970,438 | 933,696 | 9806 | 34177 | PRJEB85726 | This study |
| GW1966 | *T. citrina* | 780,516 | 749,962 | 9805 | 14836 | PRJEB85726 | This study |
| GW1967 | *T. citrina* | 494,936 | 475,406 | 8761 | 10593 | PRJEB85726 | This study |
| GW1968 | *T. citrina* | 901,608 | 862,216 | 10047 | 33861 | PRJEB85726 | This study |
| GW1969 | *T. citrina* | 16,882 | 16,156 | NA | NA | PRJEB85726 | This study |
| GW1970 | *T. citrina* | 20,476 | 10,636 | NA | NA | PRJEB85726 | This study |
| GW1971 | *T. citrina* | 712,098 | 679,378 | 9781 | 28262 | PRJEB85726 | This study |
| GW1972 | *T. citrina* | 22,484 | 21,306 | NA | NA | PRJEB85726 | This study |
| GW1973 | *T. citrina* | 10,978 | 10,146 | NA | NA | PRJEB85726 | This study |
| GW1974 | *T. citrina* | 508,826 | 488,316 | 9244 | 12131 | PRJEB85726 | This study |
| GW1975 | *T. citrina* | 941,590 | 898,910 | 9723 | 17067 | PRJEB85726 | This study |
| GW1976 | *T. citrina* | 824,232 | 785,268 | 9478 | 15731 | PRJEB85726 | This study |
| GW1977 | *T. citrina* | 705,672 | 672,298 | 9952 | 13893 | PRJEB85726 | This study |
| GW1978 | *T. citrina* | 1,272,122 | 1,214,040 | 10354 | 15114 | PRJEB85726 | This study |
| GW33333 | *T. wilhelma* | 14,653,382 | 14,217,096 | 13404 | 235544 | ERR10048047 | Wörheide et al. 2024 |
| GW41675 | *T. seychellensis* | 5,495,772 | 5,334,556 | 14519 | 80329 | PRJEB60480 | Erpenbeck et al. *2025* |

**Table S3.** Species tree topologies for *Tethya* within the 95% credible set inferred by SPEEDEMON across tested collapse threshold values (ε), using a priori species assignments following the six-species hypothesis (H6; see Fig. 2). For each ε-value, topologies are ranked by posterior probability (PP) and assigned to the corresponding species delimitation hypothesis (H4, H5.2, or H6). Topologies are presented as rooted Newick strings. Results are shown for **(a)** SPEEDEMON-SNAPPER, **(b)** SPEEDEMON-StarBeast3 with site model parameters linked across loci, and **(c)** SPEEDEMON-StarBeast3 with site model parameters unlinked across loci.

| (a) **SNAPPER** |  |  |  |  |
| --- | --- | --- | --- | --- |
| **Collapse threshold (ε)** | **Hypothesis** | **Rank** | **Species Tree Topology** | **PP** |
| 0.075 | H5.2 | 1 | ((Tau^1^,Tau^2^),(Tcit^1^+Tcit^2^,(Tmel,**out**))); | 0.834 |
|  | H5.2 | 2 | ((Tau^1^,Tau^2^),((Tcit^1^+Tcit^2^,Tmel),**out**)); | 0.087 |
|  | H5.2 | 3 | ((Tau^1^,Tau^2^),((Tcit^1^+Tcit^2^,**out**),Tmel)); | 0.078 |
| 0.100 | H5.2 | 1 | ((Tau^1^,Tau^2^),(Tcit^1^+Tcit^2^,(Tmel,**out**))); | 0.831 |
|  | H5.2 | 2 | ((Tau^1^,Tau^2^),((Tcit^1^+Tcit^2^,Tmel),**out**)); | 0.086 |
|  | H5.2 | 3 | ((Tau^1^,Tau^2^),((Tcit^1^+Tcit^2^,**out**),Tmel)); | 0.078 |
| 0.125 | H4 | 1 | (Tau^1^+Tau^2^,(Tcit^1^+Tcit^2^,(Tmel,**out**))); | 0.649 |
|  | H5.2 | 2 | ((Tau^1^,Tau^2^),(Tcit^1^+Tcit^2^,(Tmel,**out**))); | 0.182 |
|  | H4 | 3 | (Tau^1^+Tau^2^,((Tcit^1^+Tcit^2^,Tmel),**out**)); | 0.070 |
|  | H4 | 4 | (Tau^1^+Tau^2^,((Tcit^1^+Tcit^2^,**out**),Tmel)); | 0.060 |
|  | H5.2 | 5 | ((Tau^1^,Tau^2^),((Tcit^1^+Tcit^2^,Tmel),**out**)); | 0.019 |
| 0.150 | H4 | 1 | (Tau^1^+Tau^2^,(Tcit^1^+Tcit^2^,(Tmel,**out**))); | 0.852 |
|  | H4 | 2 | (Tau^1^+Tau^2^,((Tcit^1^+Tcit^2^,Tmel),**out**)); | 0.089 |
|  | H4 | 3 | (Tau^1^+Tau^2^,((Tcit^1^+Tcit^2^,**out**),Tmel)); | 0.079 |
| (b) **StarBeast3 (linked)** |  |  |  |  |
| **Collapse threshold (ε)** | **Hypothesis** | **Rank** | **Species Tree Topology** | **PP** |
| 0.0001 | H6 | 1 | ((Tau^1^,Tau^2^),(((Tcit^1,^Tcit^2^),**out**),Tmel)); | 0.403 |
|  | H6 | 2 | (((Tau^1^,Tau^2^),Tmel),((Tcit^1^,Tcit^2^),**out**)); | 0.406 |
|  | H6 | 3 | (((Tau^1^,Tau^2^),((Tcit^1^,Tcit^2^),**out**)),Tmel); | 0.290 |
| 0.001 | H4 | 1 | (Tau^1^+Tau^2^,((Tcit^1^+Tcit^2^,**out**),Tmel)); | 0.421 |
|  | H4 | 2 | ((Tau^1^+Tau^2^,Tmel),(Tcit^1^+Tcit^2^,**out**)); | 0.302 |
|  | H4 | 3 | ((Tau^1^+Tau^2^,(Tcit^1^+Tcit^2^,**out**)),Tmel); | 0.270 |
|  | H5.2 | 4 | ((Tau^1^,Tau^2^),((Tcit^1^+Tcit^2^,**out**),Tmel)); | 0.003 |
| 0.005 | H4 | 1 | (Tau^1^+Tau^2^,((Tcit^1^+Tcit^2^,**out**),Tmel)); | 0.404 |
|  | H4 | 2 | ((Tau^1^+Tau^2^,Tmel),(Tcit^1^+Tcit^2^,**out**)); | 0.323 |
|  | H4 | 3 | ((Tau^1^+Tau^2^,(Tcit^1^+Tcit^2^,**out**)),Tmel); | 0.273 |
| (c) **StarBeast3 (unlinked)** |  |  |  |  |
| **Collapse threshold (ε)** | **Hypothesis** | **Rank** | **Species Tree Topology** | **PP** |
| 0.001 | H4 | 1 | ((Tau^1^+Tau^2^,Tmel),(Tcit^1^+Tcit^2^,**out**)); | 1.0 |
| Tau^1^ and Tau^2^ = the two T. aurantium sublineages; Tcit^1^ and Tcit^2^ = the two T. citrina sublineages; Tmel = T. meloni; out = outgroup (T. wilhelma and T. seychellensis). A "+" between two lineages indicates that they were collapsed into a single species. | | | | |

**Table S4.** Species tree topologies inferred by BPP A01 across clock models and prior configurations, and their posterior probabilities (PP; topologies are shown until cumulative PP ≥ 0.95 for each analysis). Initial priors: θ ~ IG(3, 0.01), τ ~ IG(3, 0.001); broad priors: θ ~ IG(3, 0.1), τ ~ IG(3, 0.01); narrow priors: θ ~ IG(3, 0.002), τ ~ IG(3, 0.0003). Analyses that did not converge are indicated with an asterisk (*).

| **Clock Model** | **Prior settings (θ, τ)** | **Topology** | **PP** |
| --- | --- | --- | --- |
| Strict | Initial | ((((A, B), C), (D, E)), F); | 0.739 |
|  |  | (((A, B), (C, (D, E))), F); | 0.236 |
|  | Broad* | (((A, B), (C, (D, E))), F); | 0.976 |
|  | Narrow | ((((A, B), C), (D, E)), F); | 0.400 |
|  |  | ((((A, B), (D, E)), C), F); | 0.400 |
|  |  | (((A, B), (C, (D, E))), F); | 0.199 |
| Relaxed | Initial | ((((A, B), (D, E)), C), F); | 1.000 |
| A = T. aurantium sp.1; B = T. aurantium sp.2; C = T. meloni; D = T. citrina sp.1; E = T. citrina sp.2; F = outgroup. | | | |

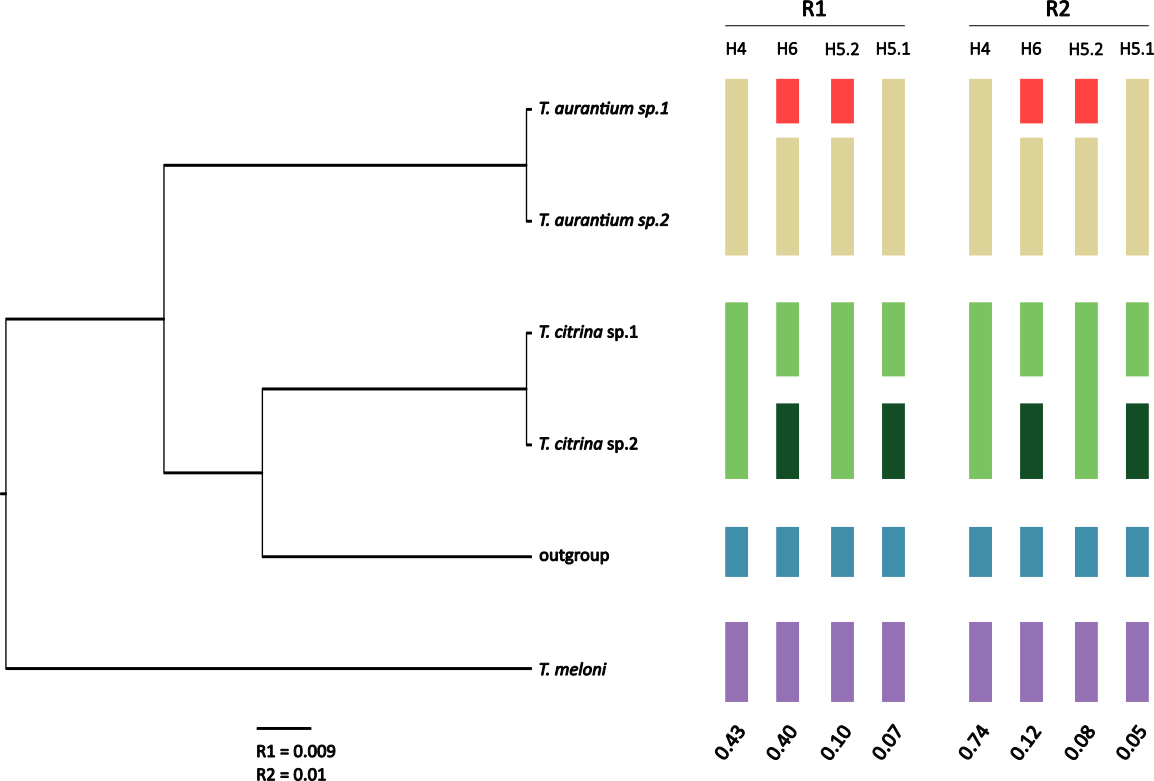

**Figure S1.** SPEEDEMON-StarBeast3 species delimitation results with a priori species assignments (H6) under a relaxed clock model and site model parameters linked across loci. Species trees are the ultrametric maximum clade credibility (MCC) trees (median node heights) for replicate analyses R1 and R2, obtained at ε = 0.001. Scale bar is shown in expected substitutions per site for both replicate analyses. Coloured bars indicate the species clusters recovered at each collapse threshold (ε); bars of the same colour denote merged lineages. Values below each column give the posterior probability (PP) of the corresponding species delimitation. Note: the analyses under the relaxed clock did not reach convergence, and the results should be interpreted as such.

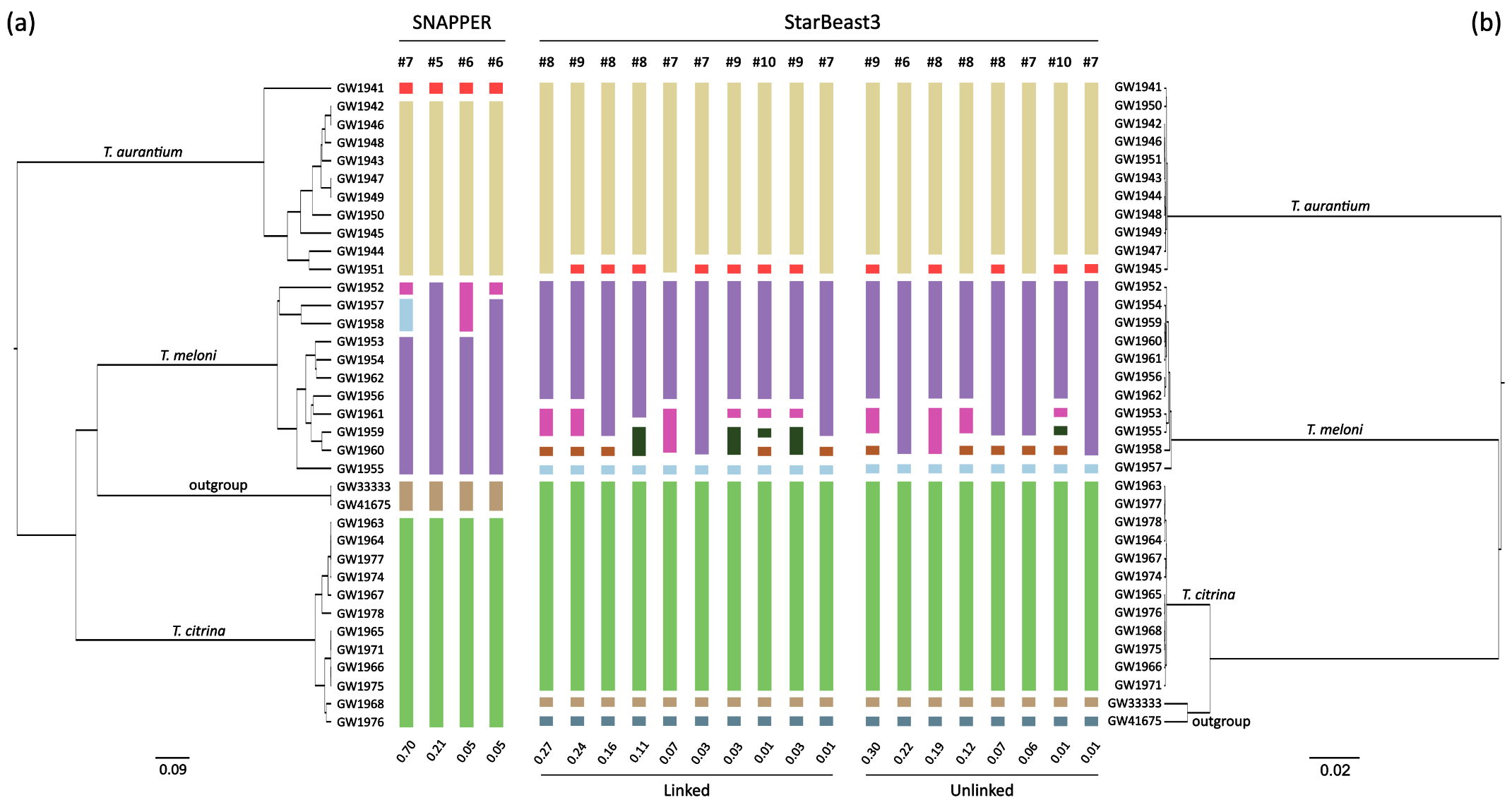

**Figure S2.** Unrestricted species discovery results obtained from SPEEDEMON under a strict clock model. For both panels, the ultrametric maximum clade credibility (MCC) species tree (median node heights) is shown as the reference topology. Coloured bars indicate the species clusters recovered within the 95% credible set. The posterior probabilities (PP) shown below each bar represent support for the corresponding species cluster assignment, not for the specific species tree topology; a given cluster count may be supported across multiple topologies. Scale bars indicate expected substitutions (mutation-scaled) per site. Only results from one of the replicate runs are shown; the second replicate run is available in the online repository (<https://github.com/PalMuc/SpongeMSCSD>). (**a**) SPEEDEMON-SNAPPER results (ε = 0.125). (**b**) SPEEDEMON-StarBeast3 results (ε = 0.001), shown for both linked (middle) and unlinked (right) site model parameters across loci. Note: the analyses shown in both panels failed to reach convergence (ESS < 200 for key parameters; see Results), and the results presented here should not be interpreted as reliable species delimitation outcomes.
